# A neural surveyor to map touch on the body

**DOI:** 10.1101/2020.06.26.173419

**Authors:** Luke E. Miller, Cécile Fabio, Malika Azaroual, Dollyane Muret, Robert J. van Beers, Ales-sandro Farnè, W. Pieter Medendorp

## Abstract

Perhaps the most recognizable sensory map in all of neuroscience is the somatosensory homunculus. Though it seems straightforward, this simple representation belies the complex link between an activation in a somatotopic map and the associated touch location on the body. Any isolated activation is spatially ambiguous without a neural decoder that can read its position within the entire map, but how this is computed by neural networks is unknown. We propose that the somatosensory system implements multilateration, a common computation used by surveying and GPS systems to localize objects. Specifically, to decode touch location on the body, multilateration estimates the relative distance between the afferent input and the boundaries of a body part (e.g., the joints of a limb). We show that a simple feedforward neural network, which captures several fundamental receptive field properties of cortical somatosensory neurons, can implement a Bayes-optimal multilateral computation. Simulations demonstrated that this decoder produced a pattern of localization variability between two boundaries that was unique to multilateration. Finally, we identify this computational signature of multilateration in actual psychophysical experiments, suggesting that it is a candidate computational mechanism underlying tactile localization.

## Introduction

In the 18^th^ century, surveyors in France completed the world’s first topographically accurate map of an entire country. To do so, they relied on the computation of *triangulation*; given a precisely known distance between two baseline landmarks, the location of a third landmark could be computed from its angles of intersection with the baseline landmarks. Countries could utilize this simple geometric computation to accurately map the location of all landmarks in their borders (Figure 1A). This is also possible using *multilateration* (or *trilateration*, more specifically) where the known distance between multiple baseline landmarks is used to compute the location of another landmark. These computations are simple yet robust ways to localize objects and therefore still used in modern surveying and global position systems.

**Figure 1.**
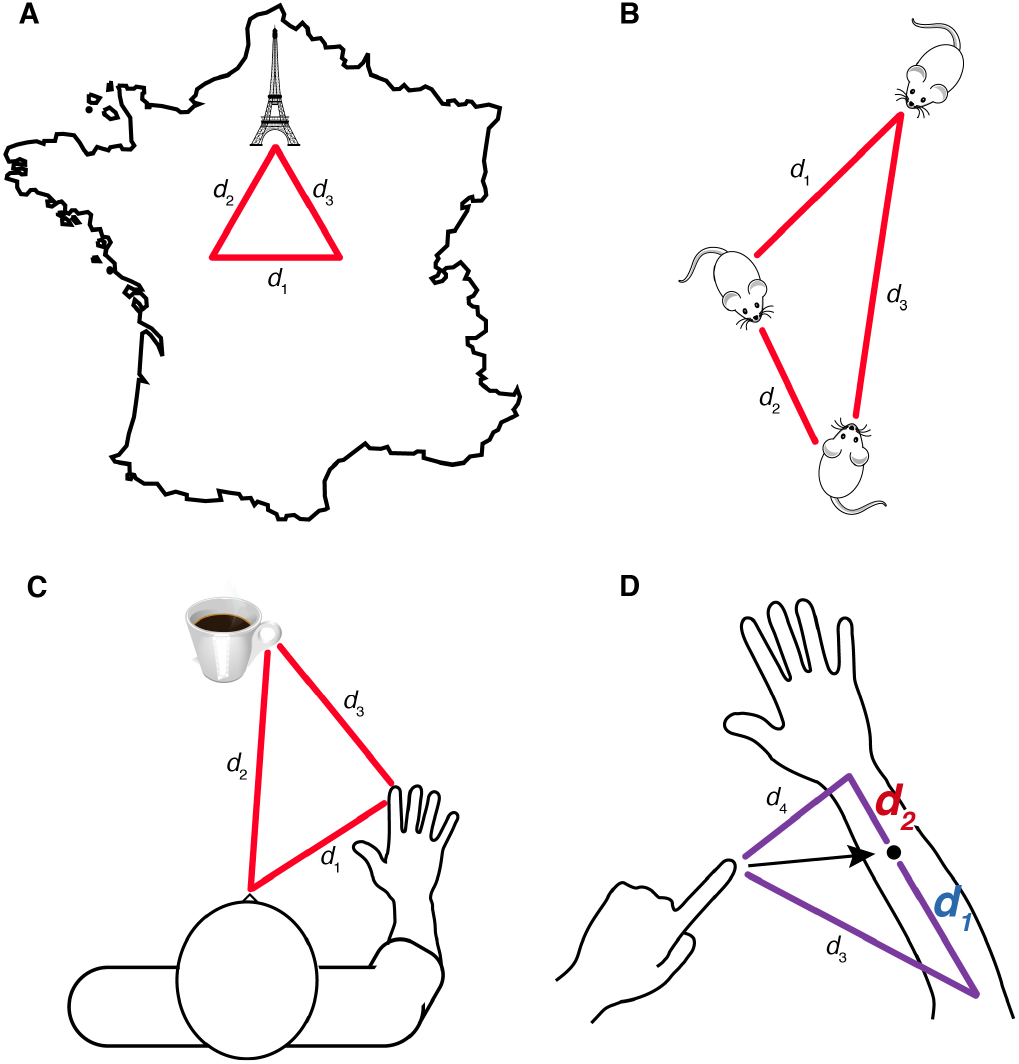
Examples of geometric computations. (A) Idealized example of how multi/trilateration can be used to localize an object within a country. Given the distance two baseline landmarks whose locations are known (*d_1_*), an object (e.g., the Eiffel tower) can be localized by calculating its distance from each landmark individually (*d_2_* and *d_3_*). (B) Path integration: By computing over distances and angles traveled (*d_1_* and *d_2_*) a rat can calculate how much it needs to travel (*d_3_*) to return to its starting position. (C) Visuomotor reaching: The distance between the hand and object (*d_3_*) can be computed by evaluating over the eye-centered hand distance (*d_1_*) and object distance (*d_2_*). (D) Egocentric tactile localization: The vector (black arrow) for reaching to a touch (black dot) on the arm may be computed using trilateration. First, an arm-centered touch location is trilaterated (see next section; Figure 2A) by computing distance between the touch and the elbow (*d_1_*) and wrist (*d_2_*). An egocentric representation of touch could be derived by further taking into the distance between the reaching hand and the elbow (*d_3_*) and wrist (*d_4_*).

Geometric computations involving manipulating distances and angles are also employed by the nervous system of animals to localize and interact with objects in the environment. During spatial navigation (Figure 1B) mammals can readily return to their starting location by taking into account all computed distances and heading directions travelled (1), a phenomenon known as path integration. Reaching to grasp a visible target is another behavior involving geometric computations (Figure 1C). To do so, the brain must compute a reach vector from distances derived from hand and target position signals (2, 3), involving transformations that take place in the frontal and parietal cortices (4, 5).

Equally crucial to localizing objects in the environment is localizing objects on the personal space of the body. In the 180 years since Weber’s seminal investigations on the sense of touch (6), researchers have extensively characterized tactile localization at both the behavioral (7–18) and neural (19–25) level. However, despite this progress, the computations underlying tactile localization remain largely unknown. Recent accounts have suggested that tactile localization requires two computational steps (26, 27). First, afferent input must be localized within a topographic map in somatosensory cortex (19). However, an activation within this map is not sufficient for localization since it alone is meaningless without a reference frame that is centered on the body. Localizing touch on the body therefore requires a neural decoder (28) that can “read” the topographic landscape of the population response within the map (22) and reference this information to stored spatial representations of the body (29). However, given that the nature of these computations—and how they might be implemented by neural circuits—remains largely unknown, it is unclear whether the brain uses geometric computations to localize objects touching the body.

We propose that, like a surveyor, the human brain employs *multi/trilateration* to localize an object in body-centered coordinates (Figure 1D). To do so, this ‘neural surveyor’ uses simple arithmetic to calculate the relative distance between the location of the afferent input and the boundaries of the body part (the baseline landmarks; e.g., elbow and wrist of the forearm in Figure 2A). This body-centered coordinate system is tied to a neural architecture that represents the body’s geometry (29) and is influenced by proprioceptive feedback about limb size (30). In the present study, we provide multiple lines of evidence that the brain may use multi/trilateration to localize touch on a limb. We first develop a Bayesian formulation of it in the nervous system. We then develop a population-based neural network model that implements trilateration, thus allowing us to identify its computational signatures. Simulations revealed that trilateration produces a unique pattern of localization variability across a limb. Finally, we identify this pattern in actual psychophysical experiments and provide further predictions of the model to be tested in future work.

**Figure 2.**
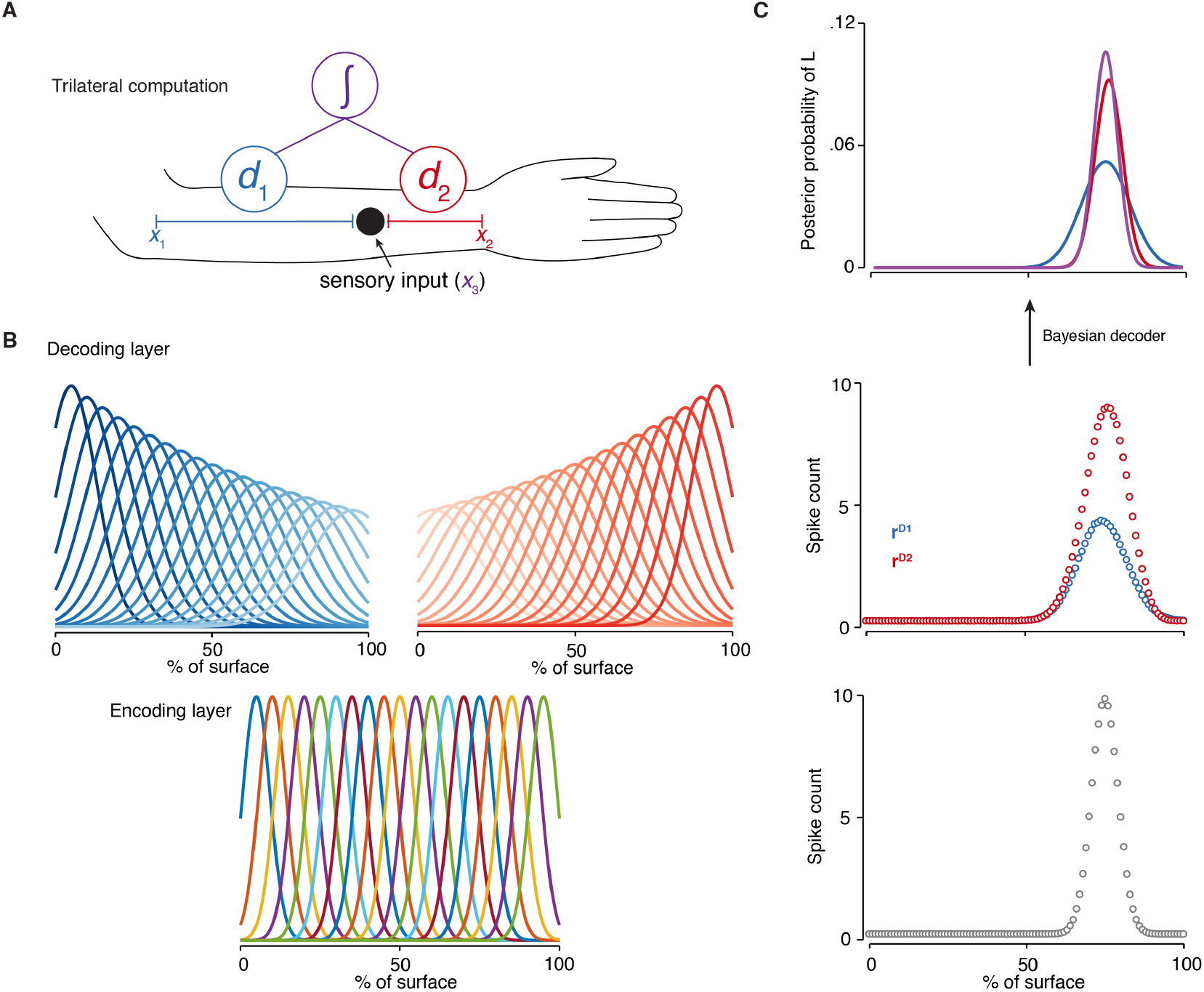
Neural network implementing trilateration. (A) Trilateral computation for tactile localization: The location of touch on the arm is computed by integrating two estimates (*d_1_* and *d_2_*) of the distance between each joint (*x_1_* and *x_2_*; elbow and wrist) and the sensory input x_3_. (B) Neural network implementation of trilateration: (lower panel) the encoding layer is composed of homogenous tuning curves across the space of the sensory surface (in units of %), (upper panel) the decoding layer is composed of two subpopulations of neurons with distance-dependent gradients in tuning properties (shown: firing rate and tuning width). The distance of a tuning curve from its “anchor” is coded by the luminance, with darker colors corresponding to neurons that are closer to the limb boundary. (C) Activations for each layer of the network averaged over 5000 simulations. Each circle corresponds to a unit of the neural network. (lower panel) encoding layer; (middle panel) decoding layer; (upper panel) posterior probabilities of localization for each decoding subpopulation (blue and red) and their integration by the Bayesian decoder (purple).

## Results

### A Bayesian formulation of trilateration

In multilateration, the distances between known locations are used to compute an unknown location. In the present paper we will focus mainly on *trilateration*, which requires calculating the distance between at least three unique locations in a common coordinate system. For simplicity, let us initially consider only a single dimension *x* of a body part (Figure 2A). Distance estimation then amounts to subtracting each location from one another:

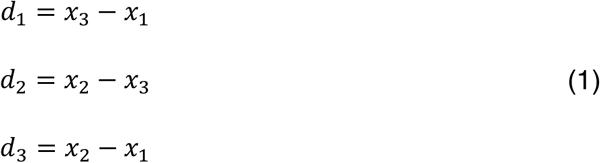

where *x*_2_ > *x*_3_ > *x*_1_, *d*_3_ is the distance between two known locations (*x*_1_ and *x*_2_) and serves as a baseline for calculating the location of *x*_3_, which is not known.

When applied to estimating the location *L* of a point of touch *x*_3_ on the limb (Figure 2A), the baseline *d*_3_ corresponds to an internal representation of limb size (29), and *x*_1_ and *x*_2_ are the boundaries of a limb-centered coordinate system. For many limbs (e.g., the forearm), these boundaries—or landmarks—are represented by the position of the proximal and distal joints; however, this is not the case for every body part (e.g., the fingertip). Given peripheral input from mechanoreceptors, *d*_1_ and *d*_2_ can be measured via a neural surveyor that ‘reads’ a population response in a central somatotopic map. Assuming noiseless point estimates (e.g., *x*_1_ = 0, *x*_2_ = 100, *x*_3_ = 75) and decoding computations, we can re-write Equation 1 to produce two estimates of location:

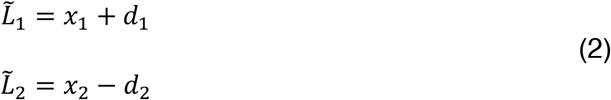

Because these estimates are defined within the same limb-centered coordinate system, a final estimate of location can be derived by taking their average, though in the case of noiseless signals both point estimates are equal and therefore redundant (i.e., both 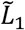 and 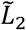 equal 75). In the nervous system, however, noise is ubiquitous (31). Sensory encoding is corrupted by receptor noise (32) which is compounded by computational operations performed by the nervous system (33). One solution for dealing with uncertainty is to take it into account when making estimates and decisions, as formalized by Bayes’ theorem (see below). If the brain does indeed take this approach, we would expect its internal estimates of the body and world are not single points but rather probability distributions over possible states.

Here we consider a probabilistic version of trilateration (Equations 1–2). In this model, an internal estimate of location is not taken as a point estimate but rather as a Gaussian probability distribution of locations. The locations along dimension *x* are therefore specified as Gaussian random variables and the landmark-centered estimates 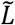 are best approximated as independent Gaussian likelihoods with distinct means and variances. Following Bayes’ theorem, touch location *L* given estimate 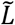—that is, the posterior distribution 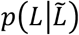—relates to:

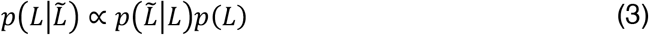

in which 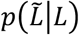 denotes the likelihood, representing probability density of the estimate 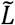 given the true location *L*, and *p*(*L*) represents prior information about the location. If we assume the prior over *L* is flat, the integrated posterior 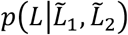 is proportional to the product the two independent likelihood functions:

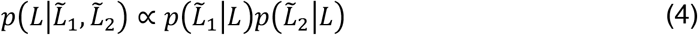

If the likelihoods are Gaussian distributions, the mean (*μ_INT_*) and variance 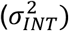 of the integrated limb-centered posterior distribution depend on the means (*μ*_1_ and *μ*_2_) variances (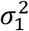 and 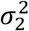) of the individual estimates:

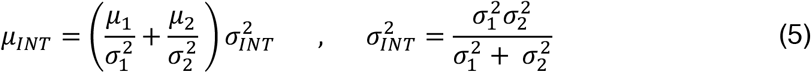

The integrated posterior thus reflects the maximum-likelihood estimate of touch location *L*, whose integrated variance is always smaller than the variance of either individual estimate. Bayesian inference of this form has been demonstrated in a range of behaviors, such as visual object recognition (34), multisensory integration (35), sensorimotor learning (36), and coordinate transformations (33, 37).

Trilateration, as formulated above, provides a computational mechanism to localize touch in body-centered coordinates. In the next section, we explore how it could be implemented by the nervous system. We describe a simple population-based neural network model that can trilaterate the location of touch from the population activity within a somatotopic map (Figure 2a). Importantly, determining the neural signatures of a trilateral computation for tactile localization will allow us to make predictions that can be validated using actual psychophysical data.

### Neural network implementation of trilateration

In the current section, we aim to ground the trilateral computation, as described in Equations 1–4, in a biologically plausible feedforward neural network. Here we implement trilateration in a probabilistic population coding network with three processing stages: First, touch is initially encoded in the activation of an early somatotopic map (encoding layer). Second, this activation pattern is transformed into body-centered coordinates (cf. Eqns. 1–2) via two decoding subpopulations whose units are tuned to distance from each landmark (cf. Eq. 2). The population activity of each decoding subpopulations reflects the likelihoods in Equation 3 (38). Lastly, the final body-centered location estimate is derived by a Bayesian decoder (39) that integrates the activity of both subpopulations (cf. Eq. 4).

The encoding layer reflects the initial somatotopic map and is composed of evenly-spaced, broadly-tuned artificial neurons with Gaussian (bell-shaped) tuning curves *f^E^* (Figure 2B; see Eq. 10 in Methods), with likelihood functions 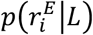 denoting the probability that location *L* caused 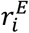 spikes in encoding neuron *i*. The likelihood function 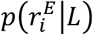 can be modeled as a Poisson probability distribution with equal mean and variance (i.e., a Fano factor of one), according to:

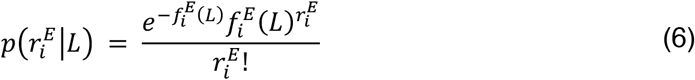

in which 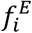 is the tuning curve of neuron *i*. The population response of the encoding neurons is denoted by a vector 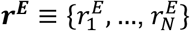, where 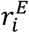 is the spike count of neuron *i*. The encoding layer thus approximates the receptive field properties of cutaneous neurons in layer IV of Area 3b whose tuning curves are relatively homogenous (40) and whose Fano factor is close to one (41).

The function of the decoding layer is to estimate the location of *L* in body-centered coordinates given the population response ***r*^*E*^** in the somatotopic map. We implemented this computation in two independent decoding subpopulations, each of which was “anchored” to one of the boundaries of the limb. The population activity ***r*^*D*^** of each subpopulation corresponds to: 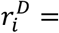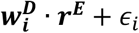 where 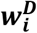 is the vector of synaptic weights connecting neuron *i* to the encoding layer, ‘∙’ is the dot product, and *ε_i_* is the additional (i.e., uninherited) Poisson noise in the decoding neuron’s spiking behavior (Eq. 6).

To embody the distance computations in Equations 1–2, the gain (i.e., peak firing rate; see Eq. 11 in Methods) of each subpopulation’s units *f^D^* formed a distance-dependent gradient (close-to-far: high-to-low gain) across the length of the limb (Figure 2B). The width of each tuning curve can be uniform in either linear or log space. In the latter case (see Eq. 12 in Methods), tuning width also forms a distance-dependent gradient (close-to-far: narrow-to-wide tuning) in linear space (42), consistent with the Weber-Fechner law. This tuning scheme has been observed in boundary vector cells (43), neurons with an analogous distance-computing function that are important for place field formation (44).

Crucially, when the neuronal noise is Poisson-like (as in Eq. 6), the gain of a neural population response reflects the precision (i.e., inverse variance) of its estimate (39). Therefore, given the aforementioned distance-dependent gradient in gain, noise in each subpopulation’s location estimate (that is, its uncertainty) will increase as a function of distance. For example, when localizing touch near the wrist, the location estimate made by the elbow-based subpopulation will be highly noisy whereas the location estimate made by the wrist-based subpopulation will be more precise (Figure 2C). A similar formulation would be appropriate for many other body parts, such as when localizing touch near the fingertip.

The exact form of the Bayesian decoder depends upon how a population response encodes the probability distribution over a stimulus. If, as several authors have argued (38, 39), population responses encode log probabilities, we can rewrite Equation 3 as follows to correspond to the maximum likelihood estimates of each subpopulation:

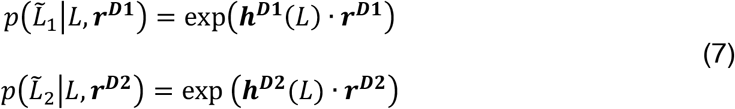

where ***h*^*D*^** is a kernel and ***r*^*D*^** is the subpopulation response. When neural responses are characterized by independent Poisson noise (Eq. 6), ***h*^*D*^** is equivalent to the log of each subpopulation’s tuning curve ***f*^*D*^** at value *L* (38, 39). Assuming that the population response reflects log probabilities, optimally integrating both estimates (Eq. 4) amounts to simply summing the activity of each subpopulation.

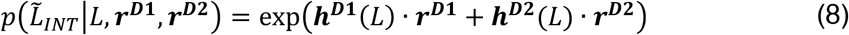

where the optimal estimate 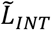 on a given trial *n* can be written as the location for which the log-likelihood of the summed population responses is maximal.

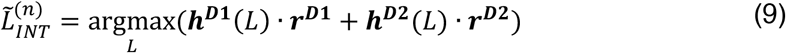

It is important to note that the integration of location estimates in Equations 4–5 assumes that they have independent sources of noise. However, given that both decoding layers are connected to the same encoding layer, they will both inherit its noise and may therefore be correlated. Simulations found that the correlation of the noise in each decoding subpopulation for each location of touch was minimal (all *r*<0.1) and can therefore be ignored. As can be seen in Figure 2C, the output of the trial-specific output of the Bayesian decoder laid out in Equations 8–9 produces an estimate whose noise is lower than both landmark-specific estimates and is consistent with maximum likelihood integration. The above formulation of trilateration is not restricted to our one-dimensional simplification, but can be generalized to the two-dimensional case as well (see Supplementary Information).

### Simulations identify a plausible neural signature of trilateration

So far we have provided a Bayesian formulation of trilateration and presented a plausible model of how this computation could be implemented in a simple feedforward network. We next investigated the localization behavior of this model by simulating single points of touch at each position within a limb-centered coordinate system (5000 simulations per location). The parameters of the units in our initial simulations were within the range of known properties of somatosensory neurons (see Methods). The units in the encoding layer had uniform gain and tuning curve width. The units in the decoding layer had distance-dependent gradients in the gain, with uniform tuning width in either linear or log space (Figure 2B). These initial simulations included only two landmarks and therefore best reflect body parts with two boundaries, such as the forearm. We consider potentially more complicated body parts, such as the fingers, later in the present report.

Both subpopulations in the decoding layer (Figure 2B) were able to localize touch with minimal constant error (Figure 3A, upper panel), demonstrating that each could produce unbiased estimates of location from the sensory input. However, as predicted given the gradient in gain, the noise in their estimates rapidly increased as a function of distance from each landmark (Figure 3B, upper panel). The pattern of location-dependent noise for each landmark-specific subpopulation was almost completely anti-correlated (*r*=–.99), forming an X-shaped pattern across the surface of the limb. Noise thus renders the estimate of each subpopulation unreliable for most of the limb.

**Figure 3.**
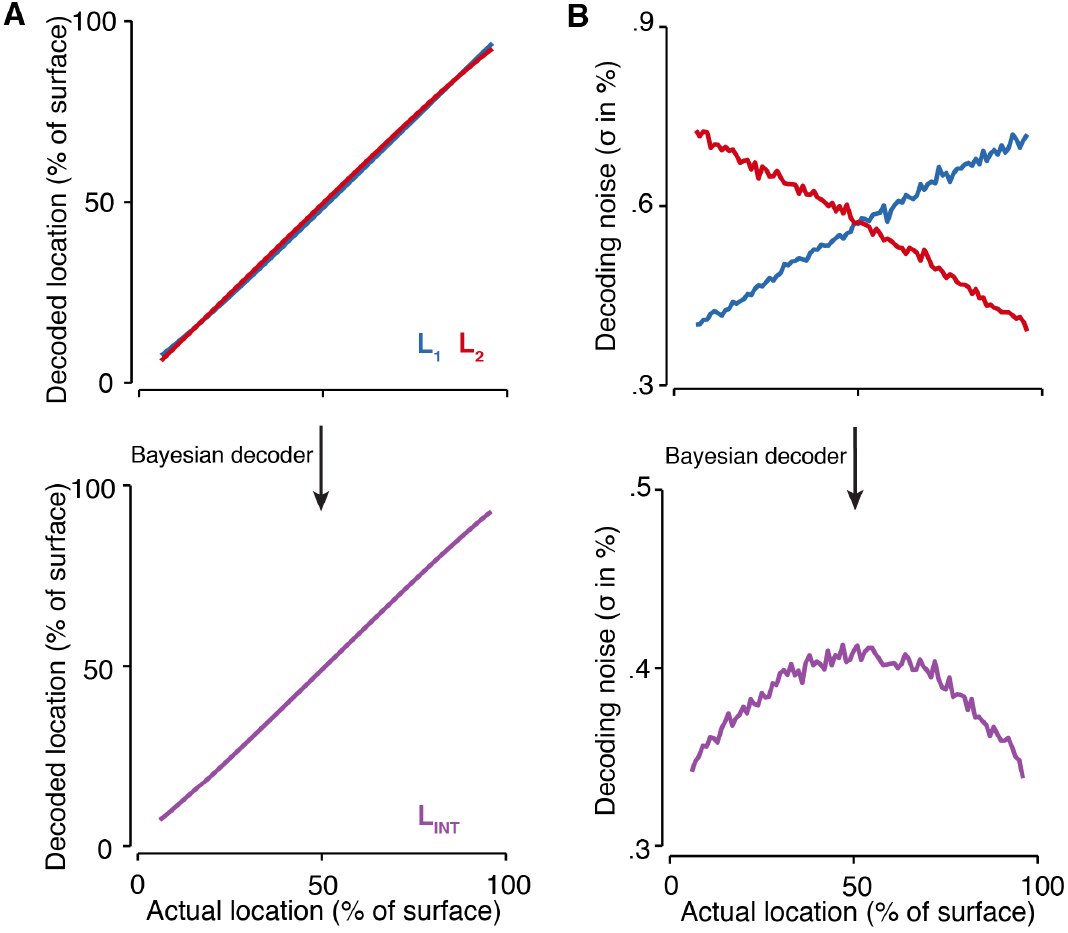
Simulation results for trilateration. (A) Localization accuracy for the estimates of each decoding subpopulation (upper panel; L1, blue; L2, red) and after integration by the Bayesian decoder (lower panel; LINT, purple). (B) Decoding noise for each decoding subpopulation (upper panel) increased as a function of distance from each landmark. Note that distance estimates are made from the 0% and 100% locations for the first (blue) and second (red) decoding subpopulations, respectively. Integration via the Bayesian decoder (lower panel) led to an inverted U-shaped pattern across the surface. Note the differences in the y-axis range for both panels.

We next examined the output of the Bayesian decoder from Equations 8–9 (Figure 2C). As expected, integrating both estimates increased the reliability (Figure 3B, lower panel; for accuracy: Figure 3A, lower panel) of localization. Intriguingly, the noise of the Bayesian decoder’s estimate formed an inverted U-shaped curve across the surface of the limb (Figure 3B, lower panel), with the lowest decoding noise near the landmarks and the highest decoding variance in the middle. This exact pattern of variability was also found when we computed the integration directly from the two simulated likelihoods (Eq. 4), demonstrating that our network optimally combines both estimates.

Unlike these initial simulations, the properties of tuning curves in real biological networks— including in Areas 1 and 2 (21, 22, 45)—are heterogenous. It is therefore important to determine whether the simulation results follow from unrealistic assumptions of homogeneity. We performed additional simulations where the tuning properties (gain and width) of each unit was varied slightly, creating local inhomogeneities in tuning. In all additional simulations (2500 distinct tuning configurations), decoding was accurate and the variability followed the inverted U-shaped pattern (Figure S1A). Regressions that modelled the decoding noise in each simulation as an inverted-U (see Methods) always found a high goodness-of-fit (mean±sd of *R^2^*: 0.73±0.08), replicating our original simulations across a variety of tuning configurations.

This signature of trilateration is independent of the chosen decoder (Eqns. 8–9) and is instead the product of integrating both decoding subpopulations. We observed the same inverted U-shaped pattern of variability with a winner-take-all decoder 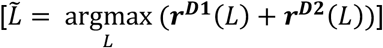, though the decoding noise was higher, consistent with its known inefficiencies. The pattern of noise is therefore a consequence of implementing trilateration of the form in our neural network and not the specific decoder used the extract information from it.

The above simulations suggest that the inverted U-shaped noise profile is a consequence of a linear probabilistic population decoder performing trilateration. However, it is possible that this decoding performance reflects inhomogeneities in the tuning of individual units in the encoding layer. Simulations rule out this possibility (Figure S1B), as our regression analysis (see above) never observed a high goodness-of-fit in any of the 2500 configurations tested (mean±sd of *R^2^*: 0.05±0.06). Importantly, there is also a conceptual argument against this possibility that bears repeating: The encoding layer merely reflects an activation within a topographic map; without the necessary extra step of referencing this activation to the body itself (as is done in trilateration), localization is impossible.

In all, our simulations suggest that inverted U-shaped pattern of variability is a computational signature of trilateration. We next conducted psychophysical experiments to test the presence of this pattern of variability in behavioral data. This is a necessary validation that our model is capturing something real in the computations underlying tactile localization. Importantly, observing an inverted U-shaped pattern of localization variability in our experiments would suggest that humans trilaterate touch location near-optimally.

### Trilateration explains tactile localization on the arm

In two psychophysical experiments, we investigated patterns of perceptual variability during tactile localization on the arm (see Methods; Figure S2). In Experiment 1 (n=11), participants localized points of touch passively applied to the volar surface of their forearm. In Experiment 2 (n=14), participants localized touch after they actively contacted an object with their forearm. Importantly, the space of possible responses was not restricted to the forearm but included both the hand and upper arm, preventing any truncation in the range of responses (Figure S10).

To assess localization, we initially fit each participant’s responses with a linear regression and the slope was taken as an overall measure of localization accuracy. Participants were generally quite accurate at localizing passive (Experiment 1; slope: 1.04, 95% CI [1.00, 1.08]; Figure 4A, upper panel; Figure S3) and active touches (Experiment 2; slope: 1.06, 95% CI [0.99, 1.12]; Figure 4B, upper panel; Figure S4). Importantly, in both experiments, we observed the expected inverted U-shaped pattern of variability (Figure 4A-B, bottom row). Thus, perceptual variability was dependent upon where the touch occurred, as predicted by trilateration. We focus on these results for the remainder of the study.

**Figure 4.**
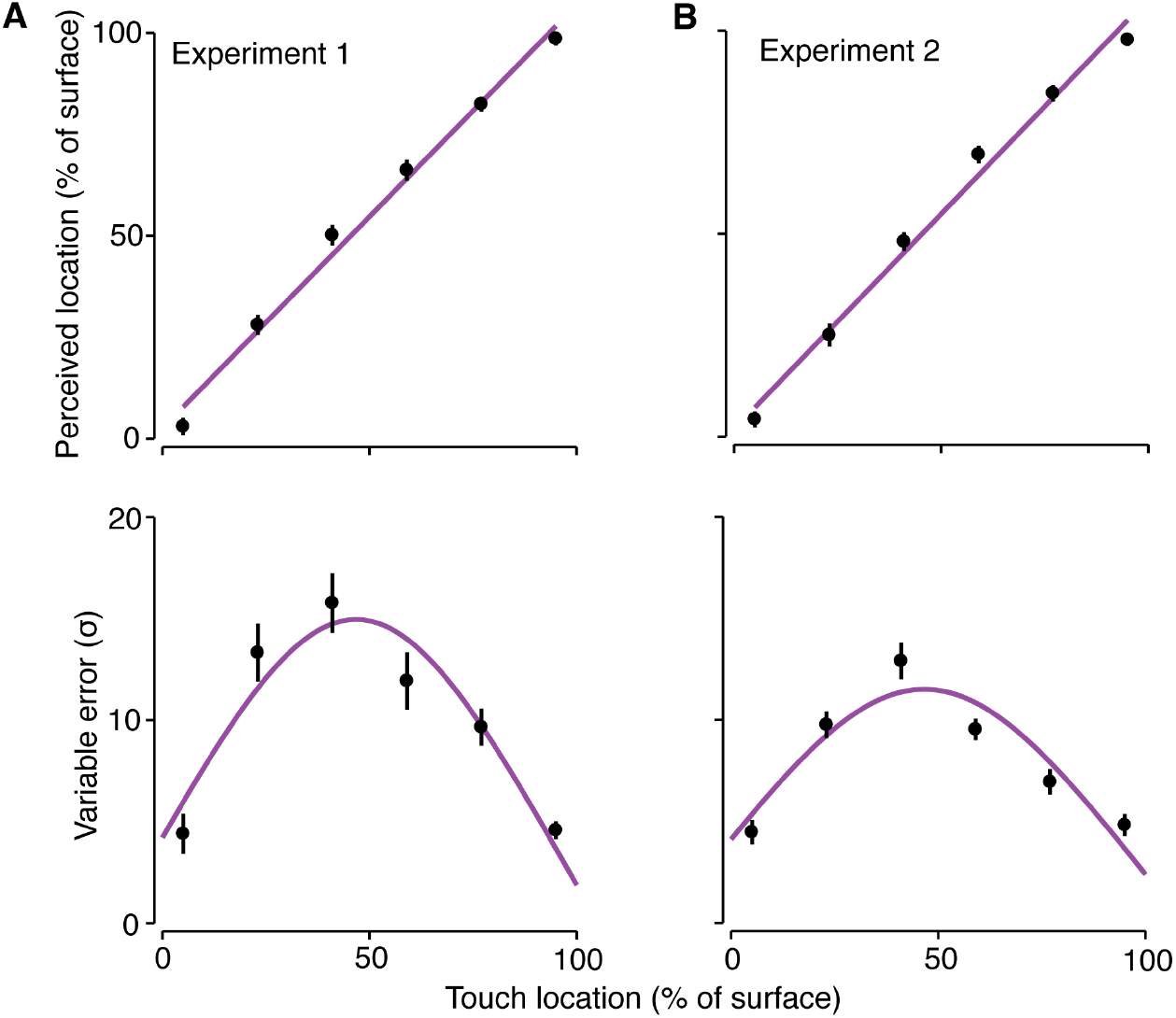
Results of behavioral experiments. The results of (A) Experiment 1 and (B) Experiment 2. Perceived location and perceptual variability as a function of touch location (0 = elbow; 100 = wrist). Tactile localization in both experiments was very accurate (upper rows). The line corresponds to a linear regression fit to the group-level data. The variable errors in units of % (lower rows) exhibited the expected signature of trilateration. The line corresponds to a trilateral regression (see below and in Methods) fit to the group-level data.

We used a reverse engineering approach (37) to validate that the observed perceptual variability was due to trilateration. Because we cannot measure the parameters of trilateration directly, we inferred them by using least-squares regression to model each participant’s variable error as a function of location (see Methods). Our regression model had three free parameters: one parameter that quantified the distance-dependent noise 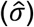 and two intercept parameters, i.e., one per landmark (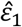 and 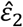). As in Equation 4, the model consisted of integrating land-mark-specific patterns of noise to form a final pattern based on an optimal estimate of location.

Trilateration explained a large portion of the location-specific patterns of variability in each experiment. We found good fits for the group-level variable errors for both passive (Figure 4A, lower panel; *R^2^*=0.89) and active touch (Figure 4B, lower panel; *R^2^*=0.86). Importantly, trilateration provided a good fit (*R^2^*>0.5) for every participant in Experiment 1 (mean±sem: 0.81±0.04; range: 0.54–0.94) and Experiment 2 (mean±sem: 0.80±0.04; range: 0.57–0.98). The fits for each participant in Experiment 1 can be seen in Figure S5 and for Experiment 2 can be seen in Figure S6.

Finally, we statistically analyzed the fit parameters for each experiment. The parameter values for each participant in Experiments 1 and 2 are listed in Table S1 and S2, respectively. We focused specifically on the noise parameter 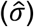, as trilateration specifically predicts that it should be greater than zero—that is, noise should scale positively with distance. The results in both experiments were consistent with this prediction (one-sample *t-*test versus zero: both *Ps*<.01, corrected). Every participant had positive distance-dependent noise (minimum of 0.27 in Experiment 1 and 0.15 in Experiment 2), demonstrating that none showed a pattern opposite from our model prediction (i.e., U-shaped variable error). This can further be observed in Supplementary Figures S5–6.

### Trilateration explains tactile localization on the finger

Thus far, we have shown that a neurally plausible implementation of trilateration accurately explains patterns of tactile localization in humans. However, these simulations and experiments were restricted to the relatively simple case of localization on body parts with two landmarks. What would trilateration look like on limbs with complicated linkage systems that would have more than two landmarks?

We simulated this situation with our neural network, which now had a third decoding population whose receptive fields were anchored midway between the other two (Figure 5A, inset). Decoding variability exhibited two hills, one on each half of the limb (Figure 5A). The presence of two hills of decoding variability was independent of the location of the third landmark and would therefore also be found in cases where the third landmark was off-center. We next explored whether this pattern of variability is observed in real localization data.

**Figure 5.**
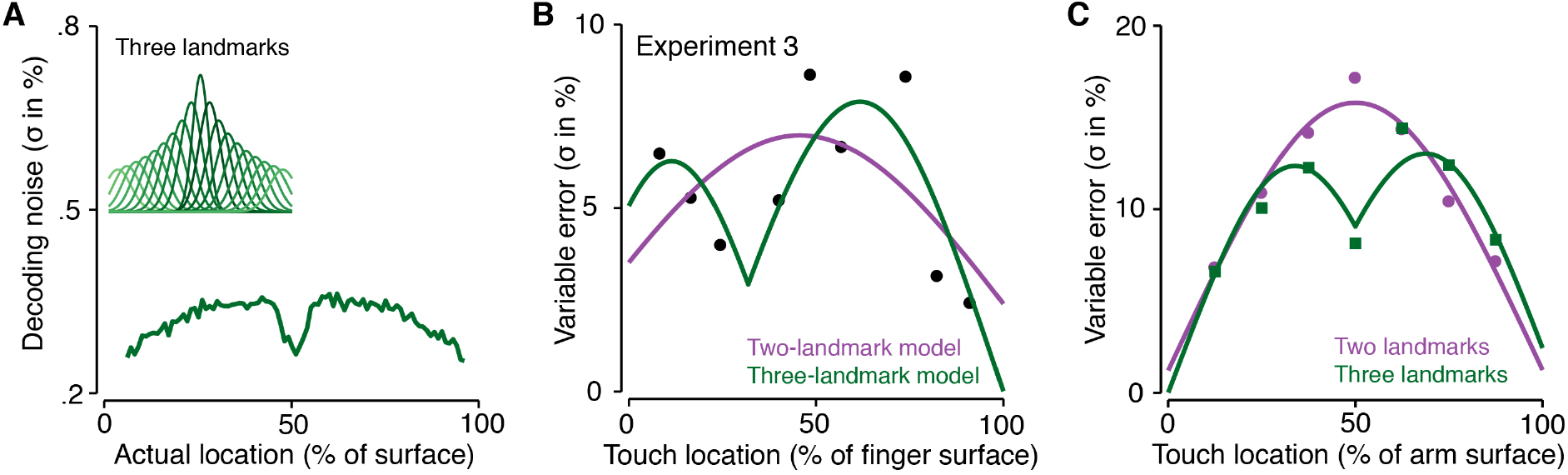
Effects of a third landmark on localization. **(A)** Simulation results of the first prediction: Adding a third landmark in the middle of sensory surface predicts an inverted W-shaped pattern of decoding variance. *Inset:* The receptive fields of the decoder subpopulation centered on this third landmark. (B) Results from a single participant in Experiment 3. The purple line corresponds to the fit of the model with only two landmarks. The green line corresponds to the fit of the model with a third landmark on the proximal interphalangeal joint. This model provides a significantly better fit. (C) Model fits to Experiment 3 from Cholewiak et al., (2003): An inverted Ushaped pattern was observed when there were two landmarks (elbow and wrist; purple); Confirming the model’s prediction, an inverted W-shaped pattern was observed when there was an additional third landmark (a stimulator) added to the middle of the forearm (green).

Using the results of these simulations as our guide, we investigated the presence of multiple landmarks within a single body part (Experiment 3). To do so, we characterized tactile localization on the index finger (n=9; ventral surface), a body part with a high degree of articulation. As expected, all participants were highly accurate at doing so (slope: 0.98, 95% CI [0.91, 1.05]; Figure S7). In the present case of the finger, four landmarks could be potentially used to trilaterate touch location: the two boundaries of the finger (metacarpophalangeal joint and fingertip) and the two intervening joints (proximal and distal interphalangeal joints). As before, each participant’s variable errors were used to explore the nature of their trilateral computation.

We fit several trilateration models related to distinct combinations of the aforementioned landmarks: a two-landmark model (i.e., boundaries only), two three-landmark models, and a four-landmark model. Trilateration provided a good fit (*R^2^*>0.5) to the location-specific patterns of variability in each participant (mean±sem: 0.73±0.04; range: 0.54–0.94; Figure S8 ; Table S3). Crucially, the data for five out of the nine participants were best fit with one of the three-landmark models (*P*<.05; Figure 5B); the proximo phalange served as the third landmark in two participants and the distal phalange in the other three. The variability for these participants were clearly characterized by two hills, as predicted by our simulations. The remaining four participants were best fit with the boundaries-only model and showed the typical inverted U-shaped pattern of variability.

These findings demonstrate that trilateration on individual body parts can include the position of intervening joints and not just body part boundaries. This raises the question about what factors influence which intervening joint is involved trilateration on the finger; this question should be explored in future work. Importantly for the purpose of the present study, all our behavioral experiments thus far have revealed broad agreement with the predictions of our population-based neural network model.

### The effect of adding a third ‘artificial’ landmark

Trilateration is not necessarily restricted to landmarks that are intrinsic to a body part (i.e., boundaries and joints). There is evidence that salient objects, such as jewelry, are represented by an internal model of a body part (46). It is therefore possible that the somatosensory system can incorporate ‘artificial landmarks’ into the trilateral computation. In these cases, we would expect the emergence of two hills of variability on a limb, one on either side of an artificial landmark that has been added to it. We explored this possibility next.

We applied our trilateration model to the results of a previously published study whose methods are conducive to addressing the above question (11). Participants in this study localized vibrotactile objects on their forearm. How these objects vibrated was varied in two conditions: (i) their vibration was at a uniform frequency across the limb; or (ii) the object at the forearm’s midpoint vibrated at a frequency that was distinct from all other locations. In the latter condition, we predicted that the midpoint object functioned as an additional landmark by which to trilaterate tactile location.

The results of our modelling confirmed these predictions (Figure 5C). As in Experiments 1 and 2, localization in the first condition was characterized by a single hill of variability, which was well-fit (*R^2^*=0.96) by a trilateration model where only the elbow and wrist were landmarks. Crucially, localization in the second condition was characterized by two hills of variability. These results were well-fit (*R^2^*=0.88) by a trilateration model where the midpoint of the forearm was an additional landmark used for localization, and bear a strong resemblance to the effect of a third internal landmark on the finger (Figure 5B; Figure S8). The somatosensory system can therefore integrate extrinsic landmarks when localizing touch on the body.

### Our results are not due to range and categorical effects

Thus far, we have argued that the observed inverted U-shaped variability profile reflects a trilateral computation underlying tactile localization. In the present section, we rule out two alternative explanations simultaneously. First, this pattern of variability may be related to a range effect (47); that is, participants were aware of the range of possible locations and therefore likelihoods at or near the boundaries were truncated. Second, and relatedly, the observed variable errors may be the result of distortions in spatial memory caused by the presence of category boundaries (48). Both explanations assume that the likelihoods for all locations 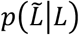 are of uniform width (i.e., *σ* does not scale with distance) but are truncated by the aforementioned cognitive processes. This truncation could theoretically produce something akin to an inverted U-shaped pattern of variability.

To rule out these alternatives, we fit the participant-level variable errors from Experiments 1– 3 with a truncation model. Like the standard trilateration model, the truncation model had three free parameters: the width of the likelihood *σ* and the locations of the truncation boundaries (see Methods). The truncation model was able to fit the data moderately well (*R^2^* mean±sem: 0.40±0.04; range: 0–0.82). However, it was a significantly worse model when directly compared to trilateration (mean±sem of ΔBIC: 7.71±0.72; range: 0.09–16.58). At the individual participant level, 32 out of 34 participants had a ΔBIC>2 (moderate evidence) and 21 of the 32 had a ΔBIC>6 (strong evidence) in favor of the trilateration model.

Further evidence against the truncation model comes from analysis of the constant errors (Figure S9). If localization responses are truncated, they would be systematically shifted inward towards the center of the body part. We did not observe this shift in any of our three experiments. Instead, we found that touch localization was slightly biased towards the wrist in Experiments 1–2 and towards the knuckle in Experiment 3. These findings are consistent with studies that have observed joint-ward biases (13, 18). Curves based upon the expected errors due to truncation could not fit the data of any participant in any our experiments (all *R^2^*<0; Figures S10–12). Furthermore, an examination of the distribution of responses made by participants (Figure S13) found that responses occasionally crossed body part boundaries—localizing touch near the wrist on the palm, for example—further ruling out the possibility that these alternative explanations could account for our experimental results.

### Computational predictions

A major benefit of our computational and neural network models is that we can use it to build predictions about how localizers should behave under different experimental conditions. We investigated three predictions made by our model of trilateration that can be tested empirically.

Correlated noise can have detrimental effects on population coding (49) and the optimality of integration (50). Our first prediction is that noise correlations should increase the magnitude of decoding noise while maintain the overall inverted U-shaped pattern. To test this, we simulated the localization of a rigid object (i.e. a continuous line), a stimulus that would introduce local noise correlations in the encoding layer. As expected, the magnitude of decoding noise (i.e., the offset of the inverted U-shaped pattern) increased as a function of object length (Figure 6A). Other manipulations that increase the magnitude of noise correlations, such as arm movements (51), would be expected to have a similar effect on decoding.

**Figure 6.**
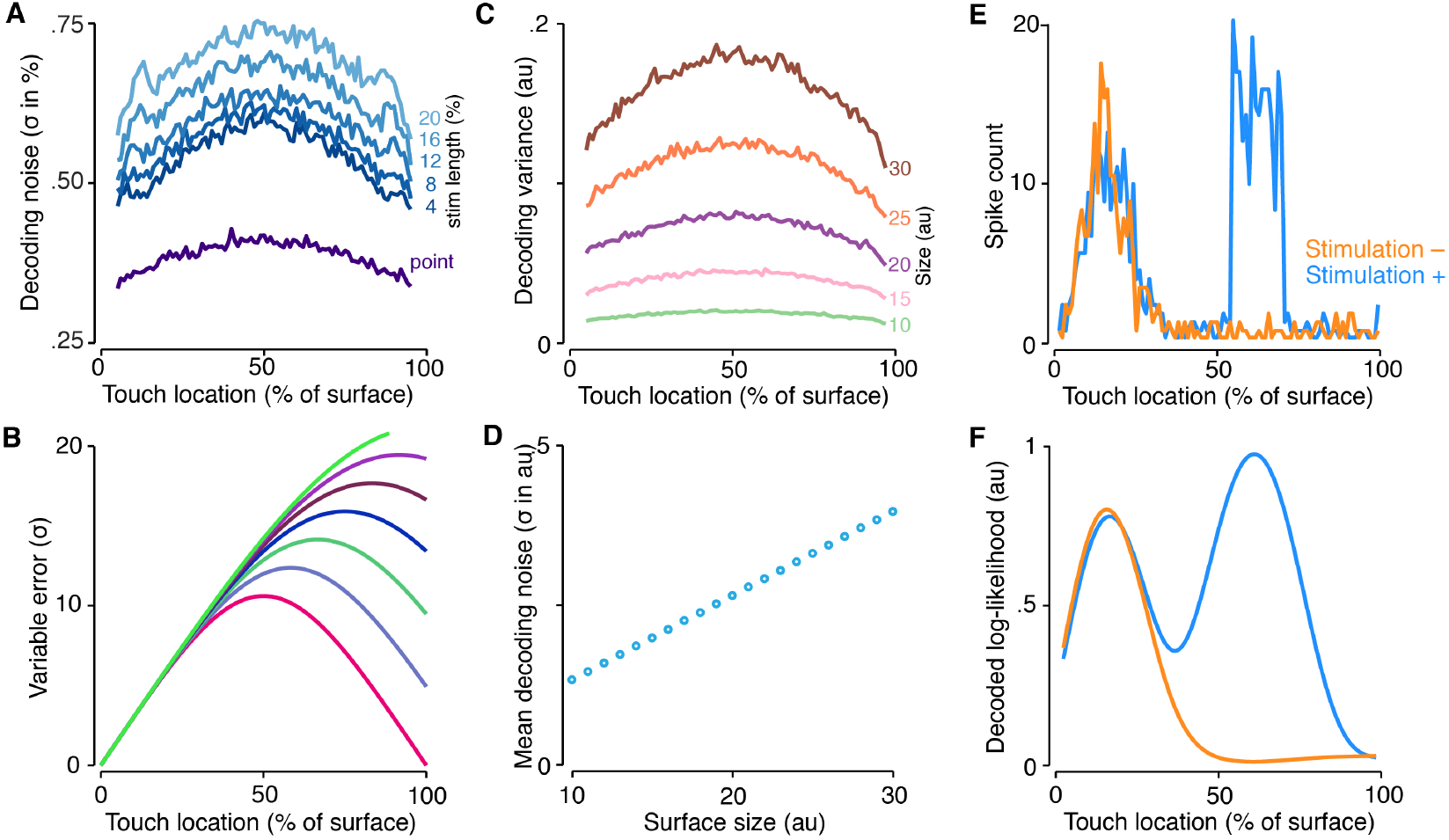
Computational and implementational model predictions. **(A)** First computational prediction: Baseline decoding noise increases as a function of stimulus length (blue curves), and is substantially higher than when the stimulus is modelled as a point (purple curve); all experiments in the present study used point-like stimuli. (B) Second computational prediction: Effect of increased variability in the second landmark location (The mark 100% on the surface). As variability increases (from 0% to 30% of tactile space; in steps of 5%), the inverted U-shaped pattern becomes more linear (less symmetrical). (C) Third computational prediction: Patterns of decoding variance for sensory surfaces of different sizes. (D) Decoding noise increases linearly as a function of size. Modifying the size of tactile space will modify perceptual variability. (E) Fourth implementational prediction: Simulated subpopulation response for touch at 15% in two conditions: without (orange curve) and with (blue curve) microstimulation of neurons coding for the third quarter of the limb. (F) Decoded log-likelihoods for these two conditions. In the case of panel E, microstimulation would modify the distance estimate derived by the Bayesian decoder.

Our second prediction is that increasing the localization variability of a single landmark will modify the shape of the perceptual variability across the limb. Specifically, as the variability of a landmark increases, the inverted U-shaped pattern of variability will become less symmetrical (Figure 6B). The variability of joint-based feedback can be modified, for example, by adding noise into the system via tendon vibration (52). In the most extreme case of completely deafferenting a joint, variability would become linear. This might not be realistic, however, given that stored offline representations of body size also play a role in the position of a landmark within a body-centered coordinate system.

Third, as our neural network is not specific to a single body part, for simplification purposes we expressed space in percentage of limb surface. However, the spatial extent of each body part is an important factor during actual tactile localization (Equation 1). Several studies have found that manipulating the internal representation of body part size (e.g., through illusions) modifies tactile spatial perception (30), including where touch is localized in space (53). Given a fixed number of units, a change in represented body part size would lead to a corresponding change in the widths of the tuning curves. Neural network simulations found that decoding noise scales linearly with a change in body part size but always maintained the inverted U-shaped profile (Figure 6C-D). Thus, localization noise should decrease when the internal representation shrinks and increase when it expands. This prediction is consistent with known changes in allocentric coding when the size of a space changes. For example, when the space of a room is elongated, place fields stretch along the axis of elongation (54), an effect mediated by boundary vector cells (44).

### Predictions for neural implementation

The computations performed by a neural network are intimately tied to the tuning properties of its units. Therefore, given the specifics of the above network’s computational architecture, we can make predictions at a neurophysiological level. Below we detail four predictions. We anticipate that they would aid in testing whether a brain region is performing multilateration of the kind embodied in our network.

First, given their importance for our implementation of multilateration, we predict the existence of distance-dependent gradients in the tuning properties of body maps in somatosensory regions. These gradients should be found in the firing rate and/or shape of the tuning curve, and follow a well-defined pattern across the representation of the body part (Figure 2B). Indeed, the wide diversity of receptive field properties present in our network (see also, Figure S15) is also found in early somatosensory regions (21). That is not to say that these gradients are a global feature that is observed in all neurons across the map. Instead, it is more likely that they will be found in neurons serving a spatial function related to localization.

Second, owing to these gradients, we predict that the range of tuning curve properties (firing rate and shape) will be higher near the border of a body part than at its midpoint. The tuning properties of units in each decoding subpopulation were almost completely anti-correlated. Therefore, neurons that belong to each distance-computing subpopulation will have tuning that is maximally different near the borders of the body part. However, since the coded distance is equal at the midpoint, the tuning of neurons with RFs near the midpoint in each sub-population should be relatively similar. Put another way, if one were to divide a body part into subsections, the extremes in response properties should be most apparent near the border.

The activity of each (sub)population in the above neural network model represents a full likelihood estimate, both its mean and its uncertainty. Each estimate is therefore encoded by a probabilistic population code (39). Our third prediction is that somatosensory neural populations will also encode the full likelihoods relevant to multilateration and will thus represent the signature distance-dependent pattern of uncertainty. Machine learning approaches, such as deep neural networks (55), could be used to decode these likelihoods from populations of neurons recorded during tactile localization. Furthermore, identifying neural responses with this pattern of population coding could determine the brain regions implementing multilateration.

Fourth, we predict that individual neurons perform specific computational functions that map onto our network; that is, encoding, distance computations, or Bayesian decoding. Since modifying the components of multilateration leads to predictable behavioral changes (see previous section), a neuron’s computational role could be identified by altering its response properties, for example, via cortical microstimulation or cooling. Stimulating one or more distance-computing neurons (Figure 6E) would have predictable effects on the shape of the likelihood encoded by the neural population (Figure 6F), influencing the decoding and therefore the behavior.

## Discussion

We proposed and tested the computation of multilateration as a candidate mechanism underlying tactile localization, directly linking sensory input to a body-centered reference frame. Neural network modeling showed that this computation can be simply implemented in a feed-forward network that integrates multiple location estimates into a single optimal surface-centered estimate. Simulations further indicated a location-dependent pattern of perceptual variability that reflects a signature of near-optimal trilateration. This signature was then found in three novel psychophysical experiment involving touch on the arm as well as one re-analyzed dataset. We conclude that multilateration is an important computation for localizing touch in the intrinsic coordinates of a sensory surface.

### Multilateration provides a unified account of tactile ‘perceptual anchors’

Tactile perception varies across the surface of an individual body part. Perhaps the most striking example is the increased perceptual precision near boundaries of joints. Evidence for this ‘perceptual anchoring’ has been observed on the arm (7, 11, 12, 16), hands (8), abdomen (56), and feet (9). Despite being first observed over 180 years ago by the psychophysicist E.H. Weber (6), the underlying reason why perception is tied to the proximity to joints is unknown. It is unlikely that ‘perceptual anchors’ have a peripheral origin since the receptive fields of mechanoreceptors are not more densely distributed near joints (57). Instead, they likely have a central explanation. For example, the integration of cutaneous and proprioceptive signals during movement may solidify joint-based landmarks as category boundaries between somatosensory representations (58). How this could be instantiated computationally has never been made explicit.

The present study suggests that the perceptual anchoring of tactile localization is a consequence of Bayesian trilateration in the nervous system. In our neural network, each decoding subpopulation is organized in reference to a specific landmark (e.g., a joint), consistent with their role as boundaries between different body-centered coordinate systems (i.e., category boundaries). Because these subpopulations represent the distance between touch and a specific landmark using a gradient of firing rates, decoding noise increases linearly as a function of distance (Eqns. 8 & 9)—the closer touch is to a landmark, the more precisely it will be decoded. Therefore, integrating estimates with distance-dependent noise naturally leads to higher perceptual precision near landmarks, both boundary-based (Experiments 1–3) and artificial (Experiment 4). Our findings thus provide a unified computational explanation of perceptual anchoring in touch and may extend to other tactile spatial phenomena (see Supplementary Information).

### Neural candidates for the distance computations

We have contrasted the process of activating a region within the somatosensory homunculus with the mapping of touch in a body-centered reference frame. The latter requires calculating the distance of the somatotopic activation—presumably in Area 3b—from the boundaries of a limb. This raises the question of which neural region(s) implements the distance computation (Equations 1–2).

Later stages of primary somatosensory processing likely implement this computation. According to our network, gradients in tuning gain and width reflect the coding of distance from a boundary. Consistent with this, a high degree of heterogeneity in firing rates has been observed in Areas 3b, 1, and 2 (21, 22, 41, 45). Furthermore, receptive field sizes of cutaneous neurons in somatosensory Areas 1 and 2 vary on a continuum from small to large (21) and often span one or more joint segments (59). These large receptive fields have previously been implicated in tactile localization (22) and may in fact reflect the tuning of distance in log space.

Neuroimaging studies in humans are consistent with this proposal. Several studies have implicated somatosensory cortex in body-centered tactile localization (60, 61). Furthermore, a recent study found evidence that tactile localization occurs at a stage recurrent processing among the three cutaneous regions of the primary somatosensory cortex (25). Recurrence may be important for the implementation of optimal estimation (62). This suggests higherorder processing rather than just topographic mapping. However, these studies are only suggestive as they lack the resolution necessary to determine the types of computations occurring in the underlying neural circuitry.

Another possibility is that distance is computed outside primary somatosensory cortex. Regions in the posterior parietal cortex play a role in the spatial coding of touch (24, 25, 60). These regions are likely important for referencing sensory signals to stored knowledge of body size (63), an important component of multilateration (Eqns. 1 & 2). A more speculative possibility is that tactile localization involves the hippocampal complex. As discussed above, the tuning of units in our decoding layer are consistent with boundary vector cells (43). Though research was initially almost exclusively focused on navigation, the hippocampal complex has been found to play a general role in spatial computation including mapping auditory (64), visual (65), and conceptual (66) spaces. Interestingly, cells with these properties were recently observed in the somatosensory cortex of foraging rats (67). It is thus possible that they may play a role in coding tactile space, including trilateration during tactile localization.

In the subsection ‘Predictions for neural implementation’, we provided a path forward for future neurophysiological research by detailing ways to identify trilateration in neural populations, though we were intentionally vague about the types of somatosensory neurons involved. One limitation of our model is that very little is currently known about how joint and skin information are combined in the somatosensory system. It is possible that, even if they are formed from multimodal signals during movement (see previous section), distance-dependent gradients are implemented in populations of purely cutaneous neurons. Alternatively, given that distances are often (though not always; see, Experiment 3) computed from joints, these gradients may be encoded by multimodal cutaneous-proprioceptive neurons (e.g., in Area 2). In line with our fourth implementational prediction, modifying the responses of neurons (Figure 6E-F) with different representational properties—and in distinct subregions of somatosensory cortex— could adjudicate between these possibilities.

### Neural implementation of the Bayesian decoder

In our neural network, the distance between touch and each landmark (Eqns. 1 & 2) is represented by the pooled activity of two subpopulations of decoding neurons (Eq. 8). By summing each subpopulation weighted by the log of each tuning curve (38, 39), the Bayesian decoder could estimate the location of touch near-optimally (Eq. 9). Unlike the encoding and decoding layers, we left the implementation of the Bayesian decoder largely unspecified. There are therefore several open questions about the nature of this computational step.

First, it is unclear whether the pooling of activity in each subpopulation would be implemented by single neurons or an entire neuronal population. Single neurons in area LIP (68) are known to integrate information from an entire sensory population (69). While this has never been directly demonstrated for somatosensory processing, several somatosensory regions have neurons with receptive fields covering an entire limb (21, 59, 70), suggesting that they pool across a population of tactile neurons as formulated in Equations 8 and 9. Alternatively, pooling could be implemented by an entire population of neurons (28). Indeed, it is often argued that Bayesian inference is best implemented at the population level (39), such as with basis functions with multidimensional attractors (62).

Second, it is unclear whether the Bayesian decoding would be implemented in somatosensory cortex or higher-order associative regions, such as posterior parietal cortex. Low-level sensory regions can implement Bayesian inference. For example, auditory spatial cues are optimally integrated by the owl midbrain during sound source localization (71). It is therefore possible that a subpopulation of neurons in somatosensory Areas 1 or 2 could optimally integrate signals from both decoding subpopulations. Instead, Bayesian decoding might be performed by somatosensory regions in the posterior parietal cortex (24). Most likely, Bayesian decoding during tactile localization is implemented by both feedforward signals in somatosensory cortex and feedback signals from posterior parietal cortex (25).

Furthermore, our behavioral experiments revealed a joint-ward bias in constant error that would be consistent with the presence of a prior in tactile space (Eq. 3). As our network produced unbiased estimates, it is best conceptualized as embodying the likelihood (Eq. 3–4). This raises the question about how a prior might be implemented in a neural network performing tactile localization. One possible mechanism to implicitly encode a prior is through biases in the distributions of tuning curves (72). However, other studies suggest that the likelihood and the prior are encoded in distinct brain regions (73). Future work should investigate how a prior is encoded in neural networks underlying the spatial coding of touch.

### Is multilateration a general spatial computation?

In the present section, we consider whether multilateration is a general spatial computation in the somatosensory system and in other cognitive domains. Whereas the present study was concerned with multilateration *within* a body part, it is worth considering whether it also underlies higher-level somatosensory behaviors such as reaching towards touch on the forearm (17; Figure 1D). Early theories of spatial perception argued that the perceived location of an object is derived from the orienting movements needed to act on it (74). Indeed everyday tactile localization typically involves reaching towards the tactile object; doing so requires transforming an initial forearm-centered code (computed via trilateration; Experiments 1–2) into egocentric coordinates (14) by integrating it with postural information about the elbow. How might multilateration play a role in this case?

One possibility is that knowing the elbow’s spatial position is sufficient for precisely reaching towards touch on the forearm, and thus multilateration is unnecessary. In contrast, the positions of both the elbow and wrist may be necessary to spatially align the reaching hand with the arm (Figure 1D). Multilateration would therefore also be crucial for the reaching component of tactile localization. Future research could adjudicate between these two possibilities by varying the relative distances between the reaching hand and arm joints, and characterizing how this modulates the level of variability in tactile localization. Only the multilateration account would predict that the proximity of the reaching hand to the wrist would have significant effects on the precision of localization on the arm.

If trilateration is indeed a fundamental computation underlying somatosensory localization, it should be employed regardless of the sensory surface. We have recently demonstrated that humans can accurately localize where a tool has been touched (75) and that mechanisms in somatosensory cortex for localizing touch on an arm may be re-used to localize touch on a tool (25). Furthermore, tool use leads to lasting changes in somatosensory perception (53, 76) that are likely driven by plasticity in somatosensory cortex (77). Thus, given the high-degree of flexibility in the somatosensory system, we predict that the computation of trilateration is also used to localize touch on tools. However, the specific details of its implementation—for example, the nature of the encoding layer—are likely somewhat different than the body and thus need to be fully worked out and investigated thoroughly.

Whether multilateration is involved in other forms of spatial cognition is also unclear. However, its equations map onto other known distance-based geometric computations implemented in the nervous system. Consider the example of reaching to grasp a coffee mug (Figure 1C). Per Equation 1, the magnitude of the reach vector (*d*_3_) can be computed simply by subtracting the vector between the eyes and the coffee mug (*d*_2_) from the vector between the hand and the eyes (*d*_1_). This operation is thought to be performed by neurons in posterior parietal cortex (3, 5).

Distance-dependent gradients in neural tuning could be one way to identify the presence of a multilateral computation. These gradients appear to be present in the neural coding of a body part’s peripersonal space (78, 79), a spatial representation involved in both keeping track of an objects distance from the body and in directing actions towards it. Several studies have found that the firing rate of visuo-tactile premotor and parietal neurons (80, 81) decreases as a non-linear function of the distance between a body part and an object. The shape of the non-linear function is consistent with the shape of the gradients in our network, suggesting that they share statistical properties. One possibility is that these neurons are involved in the reach computation described above, particularly in encoding *d*_3_.

As shown in our study, when distance is encoded by a population’s response gradient, noise in each location estimate scales linearly with distance (Equation 8). Given that distance is treated as a magnitude in this computation, distance-dependent noise is consistent with Weber’s law and may therefore be a general feature of multilateral computations. This suggests that patterns of distance-dependent noise in behavior can serve to identify multilateration in other domains. This appears to be the case in allocentric vision, where noise in the estimated location of an object is dependent upon its distance from landmarks in the scene (82). Further, the dominant source of error during path integration is noise that accumulates with distance travelled (83). Interesting, similar errors are found for path integration in the tactile domain (84), suggesting that they may involve multilateration as well.

### Conclusion

In sum, our results suggest that, like a surveyor, the somatosensory system employs near-optimal multilateration to localize a tactile stimulus. This computation is likely implemented, at least partially, in the somatosensory cortex. Future work should address how multilateration can be extended to cases of localization in two (see Supplementary Information) or three dimensions (15), as well as when touch occurs under more dynamic contexts (17, 18). Furthermore, it remains to be seen to what extent other spatial behaviors—such as path integration, allocentric vision, and reaching—could be reformulated as implementing multilateration.

## Acknowledgments

This work was supported by the ANR-16-CE28-0015 Developmental Tool Mastery to A.F., the IHU CeSaMe ANR-10-IBHU-0003, the ANR-19-CE37-0005 to A.F. & L.E.M and it was performed within the framework of the LABEX CORTEX (ANR-11-LABX-0042) of Université de Lyon. L.E.M. was further supported by a fellowship from the Donders Centre for Cognition.

## Author Contributions

L.E.M. and W.P.M. conceived of the computational model; L.E.M., R.B., and W.P.M., designed the neural network; L.E.M., performed the neural network simulations and model fitting; L.E.M., C.F., D.M., and A.F. conceived of the behavioral experiments; C.F., and M.A. performed the behavioral experiments; L.E.M. and C.F. analyzed the behavioral data; L.E.M., A.F. and W.P.M. wrote the original draft of the manuscript; All authors provided feedback on the manuscript and approved its final form.

## Materials & Methods

### Neural network modeling

#### Network parameters

We devised a simple two-layer feedforward neural network that implements trilateration to localize touch on a sensory surface. Each layer of the network was composed of artificial neurons whose preferred locations were evenly spaced across the sensory surface in our initial simulations (Figure 2; Table S4). The space of the surface was always modelled in terms of percentage (i.e., 0-100% of the surface). The properties of the units in each layer approximated important aspects of actual neurons found in the somatosensory cortex.

All units in the neural network were modelled as broadly tuned Gaussian tuning curves *f* of the following form:

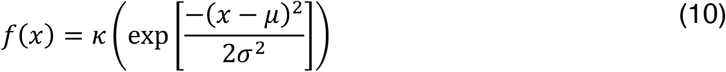

where *κ* is the peak firing rate (i.e., gain), *μ* is the tuning center, *σ* is the tuning width, and *x* is the stimulus location. As described in the Main Text, the values of *κ* and *σ* depended on the specific layer of the neural network. For tuning curves *f^E^* in the encoding layer, both their gain and width were independent of location. For tuning curves *f^D^* in the decoding layer, both the gain and width could exhibit distance-dependent gradients.

When the response properties of units exhibit independent Poisson noise—as is the case for primary somatosensory neurons—the overall gain of a population response *r* corresponds to how precisely it encodes a variable (39). As described in the Main Text, the encoding of tactile location in each decoding subpopulation exhibited distance-dependent noise. The gain of our decoding units therefore exhibited the following distance-dependent gradient:

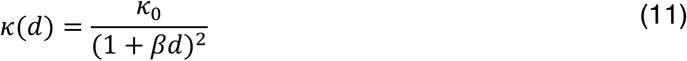

where κ_0_ corresponds to the gain of the tuning curve centered on the landmark’s location (i.e., distance zero), *d* is the distance from the center of the tuning curve (*d* ≥ 0) and the landmark, and *β* is a scaling factor.

The tuning width of units with respect to distance could either be uniform in linear or log space. In the latter case, *σ* also exhibited a distance-dependent gradient of the following form:

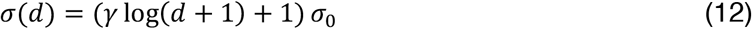

where *σ*_0_ corresponds to the width of the tuning curve centered on the landmark’s location, *d* is the distance from the center of the tuning curve and the landmark (*d* ≥ 0), and *γ* is a scaling factor.

Each unit in the decoding layer was fully connected to each unit in the encoding layer via a synaptic weight vector ***w*^*D*^**. We used the Matlab function *fmincon* to find the positive-valued weight vector that produced the decoding unit’s pre-specified tuning curve.

#### Network simulations

To investigate the consequences of a trilateral computation, we simulated 5000 instances of touch at wide range of locations (5% to 95%; see behavioral study) on the sensory surface using the above network. The values for the above parameters in the encoding and decoding layers can be seen in Table S4. Sensitivity analysis demonstrated that the pattern of results in the Main Text was not dependent on our chosen parameter values. In these initial simulations, all units of each layer shared the same parameter values. We used a maximum log-likelihood decoder to localize touch from the overall response of each subpopulation separately or added together (see Main Text). We found an identical pattern of results using a winner-take-all decoder. Results did not depend on whether tuning width in the decoding layer was uniform in linear or log space.

In follow-up simulations, we investigated the extent to which localization was influenced by the homogeneity in tuning curve parameters. We therefore simulated 2500 decoding layers where the values for *κ* and *σ* of each unit were corrupted with Gaussian noise (mean: 0, standard deviation: 0.1*parameter value). For all decoding layers, we simulated 500 instances of touch at each location. To quantify the presence of an inverted U-shaped variance profile, we fit the variance profile of the decoding with the following regression equation which produces a concave shape:

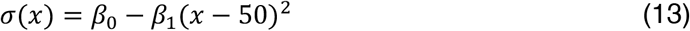

We also investigated the probability that inverted U-shaped variance was the consequence of a highly heterogenous encoding layer. We simulated 2500 encoding layers whose values for *κ* and *σ* of each unit were randomly selected from the range of values in the distance-dependent gradients of the decoding layer (Eqns. 11–12). All neural network simulations were implemented using custom code in MATLAB (MathWorks Inc.).

### Behavioral experiments

#### Subject details

Thirty-six right-handed participants in total completed our behavioral experiments. Twelve participated in Experiment 1 (8 females, 24±0.63 years of age), fifteen in Experiment 2 (9 females, 24.2±0.56 years of age), and nine in Experiment 3 (5 females, 27.9±1.30 years of age). One participant from both Experiments 1 and 2 was removed due to inability to follow task instructions. All participants had normal or corrected-to-normal vision and no history of neurological impairment. Every participant gave informed consent before the experiment. The study was approved by the ethics committee (CPP SUD EST IV, Lyon, France).

#### Tactile localization on the forearm (Experiments 1&2)

The task of participants was to localize touches applied passively (Experiment 1) or actively (Experiment 2) to their forearm, which was hidden behind an occluding board. In an experimental session, participants completed two tasks with distinct reporting methods (order counterbalanced; combined in the results of the Main Text). In the ‘drawing task’, participants indicated the corresponding location of touch on a downsized drawing of a human lower forelimb (i.e., forearm and hand). In the ‘external space task’, participants moved a cursor to indicate the corresponding location of touch within in an empty LCD screen (70 x 30 cm; white background) placed directly next to and in parallel with the arm. The location of the elbow and wrist were not indicated on the drawing or in the empty screen. Participants never received feedback about their performance.

In each experiment, there were six evenly-spaced touch locations between 5% to 95% the length of the arm (18% intervals; elbow-to-wrist; mean arm length: 23.9±0.4 cm). In each task, there were ten trials per touch location, making 60 trials per task (pseudo-random order) and 120 trials in total. See the Supplementary Information for more methodological detail.

#### Tactile localization on the finger (Experiment 3)

The task of participants was to localize touches applied passively to the ventral surface of their index finger. Participants reported the perceived location of touch on a life-sized image of their own finger (knuckle-region to tip). The image was presented against a black background on a computer screen. On each trial, the touch was presented at one of nine locations, three locations per phalanx (at 25, 50, and 75% of the actual phalanx length). Each location was touched a total of 20 times, for 180 trials in total. The specific location for each trial was chosen pseudorandomly. Participants never received feedback about their performance. See the Supplementary Information for more methodological detail.

### Statistical analyses and modelling

For Experiments 1–3, we used least-squares linear regression to analyze participants’ localization judgements and computational modelling to characterize their variable errors. To analyze Experiment 3 from Cholewiak and colleagues (2003)(11), we extracted the data from figures in their paper and then fit it with our computational model.

Our model of trilateration in the somatosensory system assumes that the perceived location of touch is a consequence of the optimal integration of multiple independent location estimates (Equations 4–5 & 8–9). Trilateration predicts that noise in each estimate varies linearly as a function of the distance of touch from a landmark *i*, the number *N* of which will vary across body parts. For any location of touch *L* along a tactile surface, the variance in a landmark-specific location estimate 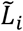 can therefore be written as follows:

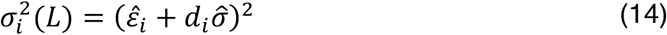

in which 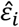 is a landmark-specific intercept term, *d*_*i*_ is the distance between touch location *L* and the landmark (Equations 1 and 2), and 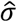 is the landmark-independent magnitude of noise per unit of distance. The distance-dependent noise for the integrated estimate takes into account the uncertainty in all *N* estimates involved in the trilateral computation on a body part.

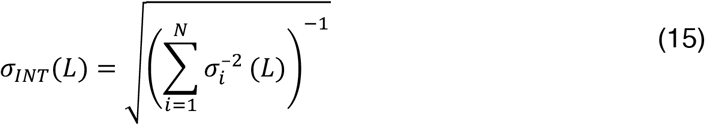

There are two landmarks for the arm (Experiments 1&2) and up to four for the finger (Experiment 3). We inferred the values of the above parameters and using a reverse engineering approach (37). All modelling for each experiment was done with the combined data from all localization tasks. Each participant’s data was also fit with a model of boundary truncation. See the Supplementary Information for more detail on our model fitting and model comparison procedures.

