## Supplementary Information for "A neural surveyor to map touch on the body"

The Supplementary information includes:

**Supplementary experimental and analytical details**

- Experiment 1: Passive touch on the forearm
- Experiment 2: Active touch with the forearm
- Experiment 3: Passive touch on the finger
- Statistical analyses and modelling
  - Localization accuracy
  - Modelling perceptual variability (Experiments 1&2)
  - Modelling perceptual variability on the index finger (Experiment 3)
  - Modelling boundary effects and model comparison
  - Modelling Experiment 3 from Cholewiak and colleagues (2003)

**Supplementary neural network and computational models**

- A two-dimensional implementation of trilateration
- Additional applications of the multilateration network architecture
- A general computational model of multilateration with *N* landmarks
- SI References

**Supplementary Tables 1–4**

**Supplementary Figures 1–18**

#### Supplementary experimental and analytical details

**Experiment 1: Passive touch on the forearm**

During the task, participants were seated comfortably in a cushioned chair with their torso aligned with the edge of a table and their right arm resting on the table top behind an occluding board. On the surface of the table, an LCD screen (70 x 30 cm) lay backside down in the length-wise orientation; the edge of the LCD screen was 5 cm from the table's edge. The center of the screen was aligned with the participant's midline.

The task of participants was to localize touches applied passively to their arm. In an experimental session, participants completed two tasks with distinct reporting methods (order counterbalanced across participants; combined in the results of the Main Text). In the 'drawing task', participants used a cursor to indicate the corresponding location of touch on a downsized drawing of a human arm (17 cm in length; elbow region to fingertips); no indication was given about the position of either the elbow or wrist. The purpose of using a downsized drawing was to dissociate it from the external space occupied by the real arm. The drawing began 15 cm from the edge of the table, was raised 5 cm above the table surface, and was oriented in parallel with their real arm. The red cursor (circle, 0.2 cm radius) was constrained to move in the center of the screen occupied by the drawing. However, participants were not constrained to only make responses on the forearm region of the drawing, thus preventing any truncation in the range of their responses. In the 'external space task', participants used a cursor to indicate the corresponding location of touch within in an empty LCD screen (white background). The cursor was constrained to move along the vertical bisection of the screen and could be moved across the entire length of the screen. It is critical to note that in this task, participants were forced to rely on proprioceptive information about their arm position as no other sensory cues were available to do so.

In each experiment, unknown to the participant, there were six evenly-spaced touch locations between 5% to 95% the length of the arm (18% intervals; elbow-to-wrist; mean arm length: 23.9±0.4 cm). In each task, there were ten trials per touch location, making 60 trials per task and 120 trials in total. The specific location for each trial was chosen pseudo-randomly. The entire experimental session took approximately 45 minutes.

The trial structure for each task was as follows: In the 'Pre-touch phase', participants sat facing the computer screen with their left hand on a trackball. A red cursor was placed at a random location within the vertical bisection of the screen. A cue (tap on the right shoulder) indicated an impending touch on the volar surface of the forearm. Touch was applied with a von Frey microfilament at a suprathreshold level of stimulation (45.3 mN of force) for approximately one second. In the 'Localization phase', participants made their task-relevant judgment with the cursor, controlled by the trackball. Participants never received feedback about their performance.

**Experiment 2: Active touch with the forearm**

The experimental procedures were identical to Experiment 1 with the following exceptions. Throughout the experiment, the participant's right elbow was placed upright in a padded support with the entire arm hidden from view behind a long occluding board. The task of participants was to localize touches that resulted from active contact between their right arm (mean arm length: 23.5±0.5 cm) and an object (rounded tip plastic cylinder; 1 mm radius). The arm was placed at a height necessary for a 1 cm separation between the object and the forearm at a posture parallel with the table. To minimize auditory cues during the task, pink noise was played continuously over noise-cancelling headphones. During each trial, a 'go' cue (tap on the right shoulder) indicated that they should actively bring their forearm to its upright posture into contact with the object, placed at one of six locations (5% to 95% of forearm length, evenly spaced). Participants were instructed to attempt to hit the object with the same speed and force across trials, though this was not measured. The number of trials and reporting methods were as in Experiment 1.

**Experiment 3: Passive touch on the finger**

During the experiment, participants were seated comfortably in a cushioned chair with their torso aligned with the edge of a table and their right arm resting on the table top behind an occluding board. The task of participants was to localize touches applied passively to the volar surface of their index finger. Touch was applied using a von Frey filament whose force (45.3 mN) was chosen to be suprathreshold for all participants. Participants reported the perceived location of touch by using a trackball to move a green cursor to the corresponding location on a life-sized image of their own finger (knuckle-region to tip). The image was presented against black background on an LCD screen. On each trial, the touch was presented at one of nine locations in total; there were three locations per phalanx (at 25, 50, and 75% of the actual phalanx length). The exact location of each touch in finger-centered coordinates thus varied slightly from participant-to-participant. Each location was touched a total of 20 times, for 180 trials in total. The specific location for each trial was chosen pseudo-randomly.

Participants never received feedback about their performance. The entire experimental session took approximately 45 minutes.

#### Statistical analyses and modelling

*Localization accuracy*

We used least-squares linear regression to analyze the localization accuracy of each task in each experiment. The mean localization judgment for each touch location was modelled as a function of actual touch location. Accuracy was assessed by comparing the group-level confidence intervals around the slope to zero and one. To standardize the data for each participant, all judgments were converted to a percentage of total surface length.

The procedure for Experiments 1&2 were as follows: For the drawing task, we converted judged locations on the drawing into percentage of drawing length. For the external space task, we converted judged locations in the space of the screen into percentage of actual arm length. In the Main Text, we collapsed localization analysis across both localization tasks for the passive and active datasets. This is because performance on the drawing and external tasks was nearly identical: passive dataset (drawing vs. external; slope: 1.03 vs. 1.06) and active dataset (drawing vs. external; slope: 1.04 vs. 1.08).

For the task in Experiment 3, we converted all judged locations in the image into percentage of actual finger length. Because every participant's finger length was slightly different, the locations of touch also varied slightly between participants. The actual locations of touch were used when modelling individual participants. The average location of touch was used when modelling the group-level data.

*Modelling perceptual variability (Experiments 1&2)*

Our model of trilateration in the somatosensory system assumes that the perceived location of touch is a consequence of the optimal integration of two independent location estimates, $\tilde{L}_1$ and $\tilde{L}_2$. This is exemplified in our Bayesian formulation of trilateration (Equations 4-5) as well as our neural network implementation (Equations 8-9). As discussed the Main Text, trilateration predicts that noise in each estimate varies linearly as a function of the distance of touch from two landmarks, corresponding to the elbow and wrist for the arm (Experiments 1 and 2). For any location of touch $L$ along a tactile surface, the variance in each landmark-specific location estimate $\tilde{L}$ can therefore be written as follows:

$$\sigma_1^2(L) = (\hat{\varepsilon}_1 + d_1\hat{\sigma})^2 \quad \text{(S1)}$$

$$\sigma_2^2(L) = (\hat{\varepsilon}_2 + d_2\hat{\sigma})^2$$

in which $\hat{\varepsilon}$ is a landmark-specific intercept term that likely corresponds to uncertainty in the location of each landmark, $d$ is the distance of touch location $L$ from the landmark (Equations 1 and 2), and $\hat{\sigma}$ is the magnitude of noise per unit of distance. Note that because the noise term $\hat{\sigma}$ likely corresponds to a general property of the underlying neural network (Equation 6-9), it is the same for each landmark. The distance-dependent noise for the integrated estimate is therefore:

$$\sigma_{INT}(L) = \sqrt{\frac{\sigma_1^2(L)\sigma_2^2(L)}{\sigma_1^2(L) + \sigma_2^2(L)}} \tag{S2}$$

The three parameters in the model ($\hat{\sigma}$, $\hat{\varepsilon}_1$, and $\hat{\varepsilon}_2$) are properties of the underlying neural processes that implement trilateration and are therefore not directly observable. They must therefore be inferred using a reverse engineering approach, where they serve as free parameters that are fit to each participant's variable errors. We simultaneously fit the three free parameters to the data using non-linear least squares regression. Optimal parameter values were obtained through maximum likelihood estimation using the MATLAB routine *fmincon*. The values of both intercept parameters ($\hat{\varepsilon}_1$ and $\hat{\varepsilon}_2$) were constrained between 0.01 and 30 and the value of the noise parameter ($\hat{\sigma}$) was constrained between -1 and 1 (units: % of surface). All modelling was done with the combined data from both localization tasks. $R^2$ values for each participant in each experiment were taken as a measure of the goodness-of-fit between the observed and predicted pattern of location-dependent noise.

*Modelling perceptual variability on the index finger (Experiment 3)*

As with the previous experiments, we assumed that the boundaries of the index finger (the metacarpophalangeal joint and fingertip) are used as landmarks. However, we also considered whether localization involves one or both of the intervening joints (proximal and distal phalanges). The variance for each of these additional landmark-specific location estimate $\tilde{L}$ can therefore be written as follows:

$$\sigma_3^2(L) = (\hat{\varepsilon}_3 + d_P\hat{\sigma})^2$$
$$\sigma_4^2(L) = (\hat{\varepsilon}_4 + d_D\hat{\sigma})^2 \tag{S3}$$

Here, $d_P$ corresponds to the distance of touch from the actual location of the participant's proximal interphalangeal joint and $d_D$ corresponds to the distance of touch from the actual location of the participant's distal interphalangeal joint. Our models were therefore specific to the actual geometry of each participant's index finger.

We fit each participant's perceptual variability with four different models. The 'boundaries-only model' modelled the variance of two estimates ($\sigma_1^2$ and $\sigma_2^2$) and was therefore identical to what was fit for Experiments 1&2 (i.e., Equations S1-2). The three additional models are as follows: The 'boundaries-plus-proximal model' and the 'boundaries-plus-distal model' also modelled the contributed variance from a third estimate, $\sigma_3^2$ and $\sigma_4^2$, respectively. The 'boundaries-plus-two model' modelled the contributed variance from both additional estimates, $\sigma_3^2$ and $\sigma_4^2$. The model-fitting procedure was identical with what we described above.

To adjudicate between which model fit each participant's data best, we performed the following procedure. We first compared the *adjusted-$R^2$* across the models. Our null hypothesis was that the ground truth model of each participant was the 'boundaries-only model'. When the winning model (i.e., highest *adjusted-$R^2$*) was one of the additional three models, we used a bootstrapping procedure to determine that its fit was not due to chance. We simulated 10,000 surrogate datasets from the 'boundaries-only model' fit (i.e., the null model). For each of the nine locations, we randomly sampled 20 datapoints from a Gaussian distribution whose standard deviation was equal to the corresponding point on the fitted curve (Equation 15). We then fit the 'variable error' of the surrogate dataset with the winning model (Equations 14–16). The 'simulated fit' was then compared to the 'actual fit'. A *p*-value was calculated as the proportion of simulations with a greater $R^2$ than the actual fit ($p$=.05 corresponds to 500 out of 10,000 simulations).

*Modelling boundary effects and model comparison*

Boundary truncation provides one alternative model to multilateration This model assumes that the estimate of location $\tilde{L}$ corresponds to a Gaussian likelihood whose variance is *identical* at all points on a body part. The inverted U-shaped variability arises because these likelihoods are truncated by a boundary, either by the range of possible responses or by a categorical boundary (e.g., between forearm and hand). We can model each likelihood $p(\tilde{L}|L)$ as a normal distribution $N(\mu_L, \sigma_L)$, where $\mu_L$ is the location of touch $L$ and $\sigma_L$ is the standard deviation. The posterior estimate $p(L|\tilde{L})$ then corresponds to a likelihood truncated at $\gamma_1$ and $\gamma_2$, where $\gamma_2 > \gamma_1$. Doing so will distort the mean and variance of the posterior estimate.

We fit this truncation model to the participant-level variable errors in each of our experiments. The standard deviation for each location, $\sigma_T(L)$, was determined by truncating a normal distribution at $\gamma_1$ and $\gamma_2$ using the *makedist* and *truncate* functions in MATLAB. The model therefore had three free parameters, $\sigma_T$, $\gamma_1$ and $\gamma_2$. The value of $\sigma_T$ was constrained between 1 and 40; $\gamma_1$ between -30 and 30; and $\gamma_2$ between 70 and 130 (units: % of surface). These ranges—particularly for $\gamma_1$ and $\gamma_2$—are quite unrealistic, but were chosen to maximize a good fit with the variable errors.

We used the Bayesian Information Criterion (BIC) to compare the boundary and multilateration models. We only considered multilateration models with two estimates in this analysis. Note that this has detrimental effects on the model fit for participants in Experiment 3 whose data was characterized by more than two estimates. The difference in the BIC (∆BIC) was used to determine a significant difference in fit; consistent with convention, the chosen cutoff for moderate evidence was a ∆BIC of 2 and the cutoff for strong evidence was a ∆BIC of 6.

*Modelling Experiment 3 from Cholewiak and colleagues (2003)*

To investigate the role of adding a third landmark to the arm, we re-analyzed Experiment 3 from Cholewiak and colleagues (2003). The full experimental details can be found in their paper; here we only present those relevant for understanding our simulations. In this experiment, participants localized a vibrotactile target stimulator (either 100 or 250 Hz) located at one of seven locations on the forearm (spacing: ~12.5% to 87.5%, by 12.5%). In a second condition, the vibrotactile stimulator that was placed at the midpoint of the arm vibrated at the opposite frequency of the rest.

To report the location of touch on a given trial, participants verbalized which of the seven target stimulators vibrated. The dependent variable was therefore the accuracy of their judgment. One downside to this approach is that it conflates accuracy and variability, as locations with very high variability (i.e., the midpoint) will have a low accuracy score. We therefore estimated the pattern of variability across the arm that would have produced their observed accuracy in each of their four conditions; the accuracy data was taken from the relevant figure using the WebPlotDigitizer tool (https://automeris.io/WebPlotDigitizer).

To convert this accuracy score into a measure of variable error, we assumed that perceived location on a given trial is drawn from a Gaussian distribution that is centered on the actual location with a certain variance. We further assumed that participants based their judgment on the activated stimulator closest to their perception. These assumptions allowed us to estimate the variable error at each location that led to the observed accuracy in each condition.

To fit the estimated variable errors in the conditions with two landmarks (i.e., elbow and wrist), we used the regression equations with three free parameters described in the previous subsection. Fitting the condition with the additional landmark (i.e., elbow, wrist, midpoint) required calculating a third variance:

$$\sigma_3^2 = (\hat{\varepsilon}_3 + d_M \hat{\sigma})^2 \tag{S4}$$

Here, $d_M$ corresponds to the distance of touch from the midpoint of the arm. This model therefore had four free parameters: $\hat{\varepsilon}_1$, $\hat{\varepsilon}_2$, $\hat{\varepsilon}_3$, and $\hat{\sigma}$. We modelled each of the three-landmark conditions separately, or with the different frequencies collapsed. Both cases produced an identical outcome and so the Main Text reports the collapsed conditions.

### Supplementary neural network and computational models

#### A two-dimensional implementation of trilateration

In the Main Text, we presented a computational and neural network model of how multilateration could be implemented within a single dimension of a body part. This is obviously an oversimplification of the real geometry of limbs, which are three-dimensional volumetric objects with complex biomechanical properties. Nevertheless, simplifying the problem to a single dimension allowed us to gain traction on how the computation could be implemented. Here we demonstrate that multilateration is not restricted to this simplified case. We show that the neural network architecture we proposed in the paper can be generalized to a surface with two dimensions.

*A computational formulation of multi-dimensional trilateration*

The computational goal of a multilateration in two dimensions is as follows: Localize a point of touch on a body part by calculating its distance from the boundaries of its surface (i.e., its 'landmarks'). It is therefore clear that this goal is independent of the number of dimensions considered in the computation. However, how this computation is achieved is dependent on the number of dimensions. For the two-dimensional case we are considering here, there are two clear differences from the one-dimensional case of the Main Text:

First, distance estimates are now calculated from four landmarks. For example, for the ventral surface of the forearm, these landmarks would correspond to the elbow, wrist, and both lateral sides. Second, and relatedly, we can no longer treat these landmarks as single points in space ($x$), but as having a two-dimensional position $(x, y)$ in a body-centered Cartesian coordinate system. For simplicity, we will ignore curvature and treat the body surface as a flat $xy$ plane. Each landmark can thus be thought of as a flat line and is therefore only point-like in a single dimension.

There are several ways that multilateration can be computed in two dimensions. Perhaps the simplest case, which we will consider here, is where each landmark is only involved in computing distance in one dimension. If we take the case of the forearm: Distances in the $x$-dimension $(d_x)$ would be calculated from the elbow and wrist, whereas distances in the $y$-dimension $(d_y)$ would be calculated from its lateral sides. Given this dimension-specificity in the distance computations, we can re-write Equation 1 of the Main Text to correspond to a specific dimension.

$$d_{x1} = x_3 - x_1$$

$$d_{x2} = x_2 - x_3 \tag{S5}$$

$$d_{x3} = x_2 - x_1$$

where $x_2 > x_3 > x_1$ and $d_x$ are the distances in the $x$-dimension. The baseline $d_{x3}$ corresponds to an internal representation of limb length, and $x_1$ and $x_2$ are the boundaries of a limb-centered coordinate system in the $x$-dimension. As before, $x_3$ corresponds to the actual location of touch $L_x$, which must be estimated using trilateration. The variables $d_{x1}$ and $d_{x2}$ are the estimated distances computed from each of these boundaries. Identical computations are also performed for the $y$ dimension, which—as detailed above—are done separately. Here, the baseline $d_{y3}$ would correspond to an internal representation of limb width. The actual location of touch can now be expressed as a two-dimensional vector, $\boldsymbol{L} = [x_3, y_3]^T = [L_x, L_y]^T$.

Given the distance estimates in each dimension (Equation S5), a neural surveyor can compute an estimate of location in each dimension.

$$\tilde{\boldsymbol{L}}_1 = \boldsymbol{\varphi}_1 + \boldsymbol{d}_1$$

$$\tilde{\boldsymbol{L}}_2 = \boldsymbol{\varphi}_2 - \boldsymbol{d}_2 \tag{S6}$$

where each estimate of location $\tilde{\boldsymbol{L}}$ corresponds to distance estimates, $\boldsymbol{d}_1 = [d_{x1}, d_{y1}]^T$ and $\boldsymbol{d}_2 = [d_{x2}, d_{y2}]^T$, made from a grouping of two landmarks, $\boldsymbol{\varphi}_1 = [x_1, y_1]^T$ and $\boldsymbol{\varphi}_2 = [x_2, y_2]^T$. Furthermore, as is clear from the Equation S6, because distances are computed separately for each dimension, landmarks can be conceptualized as lines. The essentially reduces the dimensionality of the computation down to two one-dimensional computations of the form described in the Main Text.

In line with our probabilistic interpretation of trilateration, these estimates reflect independent two-dimensional Gaussian likelihoods centered on the outputs of Equation S6. An individual estimate of touch location corresponds to the posterior $p(\boldsymbol{L}|\tilde{\boldsymbol{L}})$, where

$$p(\boldsymbol{L}|\tilde{\boldsymbol{L}}) \propto p(\tilde{\boldsymbol{L}}|\boldsymbol{L})p(\boldsymbol{L}) \tag{S7}$$

in which $(\tilde{\boldsymbol{L}}|\boldsymbol{L})$ denotes the two-dimensional likelihood representing probability density of the estimate $\tilde{\boldsymbol{L}} = [\tilde{L}_x, \tilde{L}_y]^T$ given the true location $\boldsymbol{L}$, and $p(\boldsymbol{L})$ represents prior information about the location of touch within the plane. If we again assume that the prior $p(\boldsymbol{L})$ is flat within the plane, a final location estimate can be computed by integrating both likelihoods:

$$p(\boldsymbol{L}|\tilde{\boldsymbol{L}}_1, \tilde{\boldsymbol{L}}_2) \propto p(\tilde{\boldsymbol{L}}_1|\boldsymbol{L})p(\tilde{\boldsymbol{L}}_2|\boldsymbol{L}) \tag{S8}$$

Since noise in each individual estimate increases as a function of distance from each landmark (see Results in Main Text), the pattern of noise in each integrated estimate should behave as the one-dimensional case; that is, given how we have chosen to implement trilateration, we should observe an inverted U-shaped pattern of variable error in each dimensions across the surface.

*A neural network formulation of multi-dimensional trilateration*

At the neural network level, we can implement Equations S5–S8 in a network architecture that is almost identical to the one-dimensional case. Here the network is composed of broadly tuned two-dimensional Gaussian tuning curves, with distance-dependent gradients in the decoding layer. The network's units can therefore be modelled as tuning curves $f$ of the following form:

$$f(\boldsymbol{L}) = \kappa\left(\exp\left[-\frac{1}{2}(\boldsymbol{L}-\boldsymbol{\mu})^T\Sigma^{-1}(\boldsymbol{L}-\boldsymbol{\mu})\right]\right) \tag{S9}$$

where $\kappa$ is the peak firing rate (i.e., gain), $\boldsymbol{\mu}$ is the two-dimensional tuning center, $\boldsymbol{L}$ is the two-dimensional stimulus location, and $\Sigma$ is the variance-covariance matrix of the tuning curve. We assume no covariance between each dimension.

As before, our two-dimensional network embodies the distance computations (Equation S5&6) in the tuning properties (i.e., $\kappa$ and $\Sigma$) of units in two decoding subpopulations $f^D$. Therefore, each subpopulation computes both an $x$ and $y$ location for a point of touch. That is, each calculates the distance between a point of touch and two landmarks, one for each dimension. For example, imagine a point of touch on the finger: To perform trilateration, the subpopulation $f^{D1}$ would compute its distance estimates from the base of the finger ($d_{x1}$) and the medial side ($d_{y1}$), whereas $f^{D2}$ would do so from the fingertip ($d_{x2}$) and the lateral side ($d_{y2}$). Thus, the distance-dependent tuning gradients of each unit is a multiplex of its distance from each landmark (Figure S14).

Because the distance-dependent gradients are based on each dimension independently, the computation can be conceptualized as two one-dimensional versions of trilateration. That is, the distance computations in the $x$-dimension are independent of the $y$-dimension, and vice versa. This fact is highlighted by Equation S6.

For a given subpopulation of decoding units, the width $\Sigma^D$ of their tuning curves $f^D$ had distance-dependent gradients in both the $x$ and $y$ dimensions (Figure S14), corresponding to:

$$\Sigma^D(\boldsymbol{d}) = \begin{bmatrix} \sigma_x^2(d_x) & 0 \\ 0 & \sigma_y^2(d_y) \end{bmatrix} \qquad \text{(S10)}$$

where $\sigma^2(\cdot)$ is the variance of the tuning curve in a particular dimension. As described in the Main Text, it is dependent upon the distance $d$ between the center of the unit's tuning curve and each landmark (Equation 12). As a result of these gradients, receptive field shape in this network ranged from circular to oval (Figure S15). The gain $\kappa^D$ of each tuning curve $f^D$ could also have a distance-dependent gradient (Equation 11), though only in a single dimension. The results of our simulations (see below) were independent of which dimension had the gradient in gain, or whether there was one at all.

To perform trilateration, we again implement a Bayesian decoder that assumes the population responses encode log probabilities. As in the equations in the Main Text, we can rewrite Equation S7 as follows to correspond to the maximum likelihood estimates of each subpopulation:

$$\begin{aligned} p\big(\tilde{\boldsymbol{L}}_1 \big| \boldsymbol{L}, \boldsymbol{r}^{\boldsymbol{D1}}\big) &= \exp\big(\boldsymbol{h}^{\boldsymbol{D1}}(\boldsymbol{L}) \cdot \boldsymbol{r}^{\boldsymbol{D1}}\big) \\ p\big(\tilde{\boldsymbol{L}}_2 \big| \boldsymbol{L}, \boldsymbol{r}^{\boldsymbol{D2}}\big) &= \exp\big(\boldsymbol{h}^{\boldsymbol{D2}}(\boldsymbol{L}) \cdot \boldsymbol{r}^{\boldsymbol{D2}}\big) \end{aligned} \qquad \text{(S11)}$$

where $\boldsymbol{h}^{\boldsymbol{D}}$ is the log of each subpopulation's tuning curve $\boldsymbol{f}^{\boldsymbol{D}}$ at values $\boldsymbol{L}$, and $\boldsymbol{r}^{\boldsymbol{D}}$is the subpopulation response. As in the one-dimensional case, these estimates can be optimally integrated (Equation S8) by simply summing the activity of each subpopulation.

$$p\big(\tilde{\boldsymbol{L}}_{\boldsymbol{INT}} \big| \boldsymbol{L}, \boldsymbol{r}^{\boldsymbol{D1}}, \boldsymbol{r}^{\boldsymbol{D2}}\big) = \exp\big(\boldsymbol{h}^{\boldsymbol{D1}}(\boldsymbol{L}) \cdot \boldsymbol{r}^{\boldsymbol{D1}} + \boldsymbol{h}^{\boldsymbol{D2}}(\boldsymbol{L}) \cdot \boldsymbol{r}^{\boldsymbol{D2}}\big) \qquad \text{(S12)}$$

where the two-dimensional optimal estimate $\tilde{\boldsymbol{L}}_{\boldsymbol{INT}}$ on a given trial $n$ can be written as the location that maximizes the log-likelihood of the summed population responses.

$$\tilde{\boldsymbol{L}}_{\boldsymbol{INT}}^{(n)} = \underset{\boldsymbol{L}}{\operatorname{argmax}}(\boldsymbol{h}^{\boldsymbol{D1}}(\boldsymbol{L}) \cdot \boldsymbol{r}^{\boldsymbol{D1}} + \boldsymbol{h}^{\boldsymbol{D2}}(\boldsymbol{L}) \cdot \boldsymbol{r}^{\boldsymbol{D2}}) \qquad \text{(S13)}$$

Note that Equations S11–13 above correspond to two-dimensional versions of Equations 7–9 in the Main Text but otherwise reflect the exact same operations. That is, they reflect a Bayesian decoder that implements Equations S7–8 (Equations 3–4 in Main Text). Below we will see that the output of this network is essentially identical to the one-dimensional network described in the Main Text.

*Results of neural network simulations*

We simulated the ability of this neural network to localize points of touch on a two-dimensional surface (500 simulations per location). The parameters of the tuning curves were kept the same as the simulations in the Main Text (Table S4). However, the following aspects of our simulations differed: (i) We restricted simulations to the decoding layer, as the encoding layer was unnecessary for the present purposes. The network therefore consisted of two subpopulations of decoding units and the Bayesian decoder (Equations S11–13). (ii) The spacing of each tuning curve was increased to 5%, yielding 400 units (20 x 20 grid) per subpopulation to cut down the computation time of our simulations.

We found that both subpopulations in the decoding layer were able to localize touch with minimal constant error in either dimension (Figure S16A-B), demonstrating that each could produce unbiased estimates of location from the sensory input. Consistent with our one-dimensional network, we observed a rapid linear increase in the noise in their estimates that was dependent upon the distance in each dimension. Crucially, the noise of the Bayesian decoder's estimate formed an inverted U-shaped curve across the surface of the limb in both dimensions (Figure S16C-D). The pattern in each dimension was independent of the other. We can therefore view the results of our one-dimensional network as reflecting a slice down the center of a two-dimensional network performing trilateration. Importantly, these simulations demonstrate that trilateration can successfully be performed in multiple dimensions using a very basic neural network architecture.

**Additional applications of the multilateration network architecture**

Thus far, here and in the Main Text, we have predominantly considered how the above neural network applies to tactile localization. However, it is possible that there are other spatial operations that are implemented with a similar or the same neural network architecture.

*Case 1: Tactile spatial acuity*

Spatial acuity has been linked with the size of receptive fields, including acuity in the tactile domain (1). The smaller the receptive field, the higher the spatial precision of the perception. Body maps in primary somatosensory regions display a large diversity of receptive field sizes and shapes, ranging from more circular (2) to more elongated (3). For example, receptive fields representing the hairy skin tend to have a longer proximo-distal axis compared to the medio-lateral axis. This raises the question as to how the properties of units in our neural network relate to spatial precision in general.

The distance-dependent tuning gradients of the units in our neural network (Equation S10) are consistent with the diversity of receptive fields in somatosensory regions. Even given a circular "baseline" receptive field (i.e., identical width in each axis; Equation S10) at distance zero, gradients would produce a variety of oval-shaped receptive fields (Figure S15). However, as described above, the shape of the "baseline" receptive field in several body maps is itself elongated. These tuning anisotropies are thought to underlie orientation-dependent *perceptual* anisotropies in spatial precision (4). Simulations with our neural network support this proposal. By varying the baseline shape of our receptive fields, we found that perceptual anisotropies in spatial precision are largely proportional to the tuning anisotropies (Figure S17). These differences in precision are independent of the specific effects of trilateration, suggesting that multiple computational factors likely shape tactile spatial precision. Future work should attempt to partition measures of acuity into these different factors.

*Case 2: Tactile distance perception*

Humans can perceive the distance between two objects touching their skin, a phenomenon termed *tactile distance perception*. Tactile distances may in fact be a basic somatosensory feature (5), perhaps computed early in the somatosensory cortical hierarchy (6). We are unaware of any models that formalize tactile distance estimation, though conceptual models exist (see below). Here we explore how multilateration could play an important computational role in the estimation process.

According to a prominent conceptual model—the *pixel model*—tactile distance is calculated by summing up the intervening receptive fields between two activations in a topographic map (7). The proposed summation operation is based on the finding that orientation-dependent distortions in distance perception (8) appear inversely proportional to the size of a somatosensory receptive field (RF) in a given axis. Perceptual distortions arise because a decoder fails to take the anisotropies of RF shape into account and instead assumes that receptive fields are circular (i.e., isotropic). Therefore, with less intervening RFs in the long axis, the decoder estimates that the points of touch are closer together than they really are.

How exactly the summation operation of the above form would be implemented in a neural network is left unspecified. From our perspective, summation could be viewed as computationally equivalent to the distance estimates we propose for multilateration. In our network, explicit distance estimates would only require a decoder with knowledge about the distance-dependent tuning gradients. If the characteristics of these gradients are themselves proportionally linked to receptive field shapes (see Case 1, above) and the decoder fails to take any resulting anisotropies into account (as proposed in the pixel model), its estimates will be distorted in accordance with what is observed for tactile distance estimates.

Providing a formalization and neural network implementation of tactile distance estimation is beyond the scope of this paper. However, we note two points that may be important for a multilateration-based formulation. First, given that tactile distance is concerned with the intervening space between two tactile points, it is possible that each individual point itself serves as a landmark in the computation. As we report in the Main Text, this can be the case for tactile localization (Experiment 4). Second, as with localization precision, the perceived tactile distance between two points of touch is greatest near (9) or across (10) a joint. Distance estimation likely still relies on neural computations that are tied to body part boundaries. Future work on tactile distance perception should attempt to address these issues by building a formal model.

##### A general computational model of multilateration with *N* landmarks

The formulation of trilateration above and in the Main Text is restricted to simple geometries (i.e., a line or a plane). In these cases, the landmarks refer to the boundaries of a body part's surface, which are typically either a side or a joint. In several cases, other objects or features of a body part might also play a role as a landmark. For example, previous studies have found that the navel (11) functions as a perceptual anchor (i.e., landmark) for tactile localization. In the fourth experiment of the Main Text, we demonstrated that this was the case for an additional object touching the skin. Per our neural network simulations, when there is a single additional landmark on a body surface, the variable error of tactile localization is characterized by two hills (Figure 5).

While a simple plane with four landmarks (as above) might be an adequate approximation for a single surface of some body parts (e.g., forearm, palm, finger, etc.), it would fail to capture the complexity of others. For example, localization on the face may involve multilateration from many distinct landmarks, such as the jawline, lips, nose, eyes, ears, and hairline.

Here we provide a general model of multilateration in an $xy$ plane that can be applied to a variety of body part arrangements. For this model, we provide the precision-weighted versions of probabilistic inference, which is mathematically equivalent to Equations 4–5 & S7–8. Given a hypothesized arrangement of landmarks, a pattern of localization behavior in body-centered coordinates can be estimated using the following generative model that includes an arbitrary number of $N$ independent landmark-based estimates:

$$\tilde{\boldsymbol{L}}_{\boldsymbol{INT}} = \sum_{i=1}^{N} W_i \tilde{\boldsymbol{L}}_{\boldsymbol{i}} \tag{S14}$$

where

$$W_i = diag\left(\frac{\sigma_{i,x}^{-2}}{\sum_{j=1}^{N} \sigma_{j,x}^{-2}}, \frac{\sigma_{i,y}^{-2}}{\sum_{j=1}^{N} \sigma_{j,y}^{-2}}\right) \tag{S15}$$

in which $W$ is the matrix of precision-based weight assigned to current estimate $i$ in each dimension, and $\sigma_{i,}^2$ is its distance-dependent variance (distinct for each dimension) of the following form:

$$\sigma_{i,}^2 = \left(\hat{\varepsilon}_{i,} + d_{i,}\hat{\sigma}\right)^2 \tag{S16}$$

Here, $d_{i,}$ refers to the dimension-specific distance between the $xy$ location of the landmark and the $xy$ location of the touch. When the landmark is a boundary, the above neural network implements a distance computation in one dimension only and therefore the variance in the other dimension can be thought of as infinite. Other types of landmarks could be better approximated as individual points within a body part (e.g., mouth, navel, etc.) and may thus calculate distances in both dimensions. Given all $N$ estimates, the final integrated variance corresponds to:

$$\Sigma_{INT} = \left(\sum_{i=1}^{N} \Sigma_i^{-1}\right)^{-1} \tag{S17}$$

Where $\Sigma_{INT}$ is the variance-covariance matrix of the integrated estimate. If we take the simplifying assumption of no covariance in each dimension the matrix for each individual estimate $i$ takes the following form:

$$\Sigma_i = \begin{bmatrix} \sigma_{i,x}^2 & 0 \\ 0 & \sigma_{i,y}^2 \end{bmatrix} \tag{S18}$$

This generative model would produce a pattern of variable error for any arbitrarily complex assortment of landmarks. For example, simulations for the face can be seen in Figure S18. Here, we implemented both forms of landmarks (i.e., borders and points) described above. As is evident in this figure, multiple landmarks throughout a sensory surface can lead to rather complex global patterns of variability across the body part. Future research should explore the variety of landmark types when formalizing models of tactile localization within a body part.

#### Supplementary tables and figures

| Participant | $\hat{\varepsilon}_1$ | $\hat{\varepsilon}_2$ | $\hat{\sigma}$ | $R^2$ | *RMSD* |
|---|---|---|---|---|---|
| 1 | 2.12 | 2.14 | 0.28 | 0.91 | 1.31 |
| 2 | 1.91 | 3.23 | 0.28 | 0.79 | 2.28 |
| 3 | 9.46 | 0.01 | 0.34 | 0.69 | 4.13 |
| 4 | 3.57 | 2.05 | 0.29 | 0.55 | 4.06 |
| 5 | 0.76 | 1.17 | 0.27 | 0.91 | 1.36 |
| 6 | 3.50 | 5.89 | 0.35 | 0.72 | 3.31 |
| 7 | 2.62 | 0.36 | 0.45 | 0.87 | 2.85 |
| 8 | 7.53 | 2.09 | 0.51 | 0.77 | 4.34 |
| 9 | 2.08 | 0.97 | 0.29 | 0.93 | 1.29 |
| 10 | 1.66 | 0.01 | 0.65 | 0.94 | 2.50 |
| 11 | 13.50 | 4.98 | 0.27 | 0.84 | 2.03 |
| **Mean** | 4.43±1.21 | 2.08±0.59 | 0.36±0.04 | 0.81±0.04 | 2.68±0.35 |

**Table S1. Best fit parameters for Experiment 1**

| Participant | $\hat{\varepsilon}_1$ | $\hat{\varepsilon}_2$ | $\hat{\sigma}$ | $R^2$ | *RMSD* |
|---|---|---|---|---|---|
| 1 | 4.23 | 2.68 | 0.21 | 0.65 | 2.25 |
| 2 | 2.27 | 2.81 | 0.21 | 0.87 | 1.19 |
| 3 | 4.07 | 4.64 | 0.15 | 0.57 | 1.71 |
| 4 | 5.04 | 2.69 | 0.28 | 0.96 | 0.89 |
| 5 | 2.75 | 0.01 | 0.31 | 0.67 | 3.62 |
| 6 | 3.99 | 6.06 | 0.28 | 0.92 | 1.20 |
| 7 | 4.32 | 6.21 | 0.24 | 0.85 | 1.42 |
| 8 | 6.14 | 0.01 | 0.27 | 0.80 | 2.15 |
| 9 | 1.57 | 4.84 | 0.29 | 0.98 | 0.63 |
| 10 | 1.92 | 0.41 | 0.26 | 0.73 | 2.47 |
| 11 | 9.86 | 0.01 | 0.32 | 0.87 | 2.3 |
| 12 | 4.72 | 0.01 | 0.34 | 0.63 | 5.07 |
| 13 | 6.94 | 6.39 | 0.21 | 0.96 | 0.55 |
| 14 | 2.09 | 1.90 | 0.22 | 0.78 | 1.86 |
| **Mean** | 4.28±0.61 | 2.77±0.66 | 0.26±0.01 | 0.80±0.04 | 1.95±0.33 |

**Table S2. Best fit parameters for Experiment 2**

| Participant | $\hat{\varepsilon}_1$ | $\hat{\varepsilon}_2$ | $\hat{\varepsilon}_{3P}$ | $\hat{\varepsilon}_{3D}$ | $\hat{\sigma}$ | $R^2$ | *RMSD* |
|---|---|---|---|---|---|---|---|
| 1 | 9.22 | 4.77 | 0.01 | — | 0.62 | 0.73 | 2.90 |
| 2 | 1.04 | 0.1 | — | — | 0.29 | 0.79 | 1.78 |
| 3 | 5.46 | 0.35 | — | 1.65 | 0.22 | 0.63 | 1.65 |
| 4 | 5.62 | 3.01 | 0.01 | — | 0.30 | 0.67 | 1.46 |
| 5 | 9.09 | 3.68 | — | 0.01 | 0.44 | 0.71 | 2.83 |
| 6 | 6.05 | 2.18 | — | — | 0.16 | 0.54 | 1.81 |
| 7 | 3.51 | 0.01 | — | — | 0.31 | 0.83 | 1.91 |
| 8 | 2.76 | 0.01 | — | — | 0.29 | 0.76 | 1.89 |
| 9 | 5.52 | 7.45 | — | 0.01 | 0.46 | 0.94 | 1.03 |
| **Mean** | 5.36±0.90 | 2.39±0.87 | — | — | 0.34±0.05 | 0.73±0.04 | 1.92±0.20 |

**Table S3. Best fit parameters for Experiment 3**

| | $f^E$ | $f^{D1}$ | $f^{D2}$ |
|---|---|---|---|
| $\mu$ | -40:1:140 | 0:1:140 | -40:1:100 |
| $\kappa$ or $\kappa_0$ | 25 | 25 | 25 |
| $\sigma$ or $\sigma_0$ | 4.25 | 4.25 | 4.25 |
| $\beta$ | — | 0.005 | 0.005 |
| $\gamma$ | — | 0.5 | 0.5 |

**Table S4. Initial neural network parameter values**

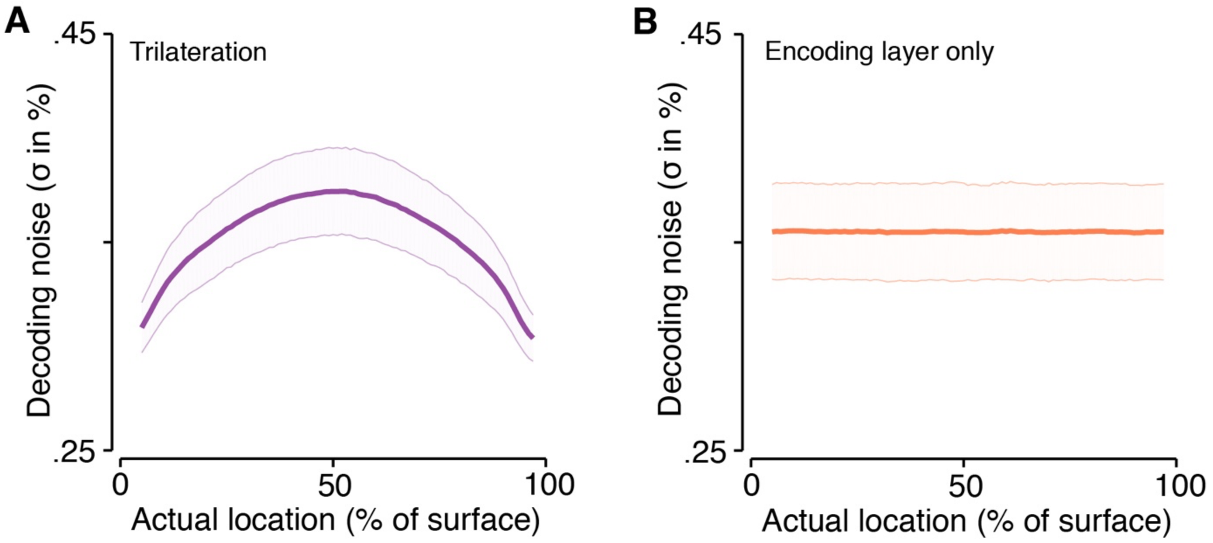

**Figure S1. Decoding  noise over a variety of parameters**

(A) The inverted U-shaped pattern of decoding noise was robust to noise in the parameter values of individual units in the decoding layer. The purple line shows the mean decoding noise over 2500 simulations and the shaded error bars correspond to the 95% confidence interval. (B) We never observed the inverted U-shaped pattern of noise when decoding from the encoding layer directly, even though the gain and tuning widths were chosen from the parameter ranges of the decoding layer (see Methods). The orange line shows the mean decoding noise over 2500 simulations and the shaded error bars correspond to the 95% confidence interval.

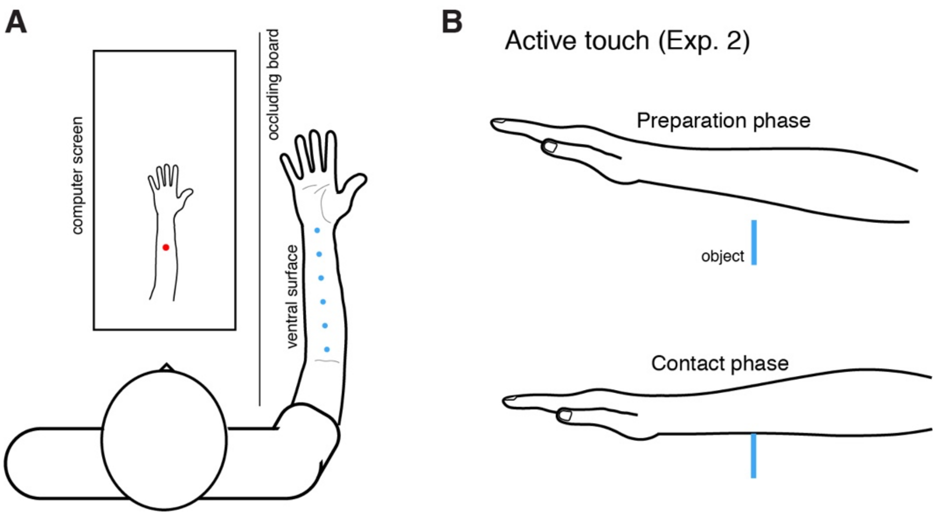

**Figure S2. Experimental setup for Experiments 1&2**

(A) Experimental setup for the localization task: Participants sat in front a computer screen (not drawn to scale) with their arm behind an occluding board. Blue circles reflect the six possible touch locations. They localized touch either on a drawing (shown) or in an empty screen (not shown). Note that the computer screen has been moved forward for illustrative purposes, but in reality was closer to the participant's torso. A similar method was used for touch on the finger in Experiment 3. (B) Participants in Experiment 2 actively contacted the object with their arm. This involved a preparation phase (top) where participants waited for a go cue. After the go cue they brought their arm downward (contact phase) and onto the object (blue line). Note that their elbow was held in place to limit the range of motion (not shown).

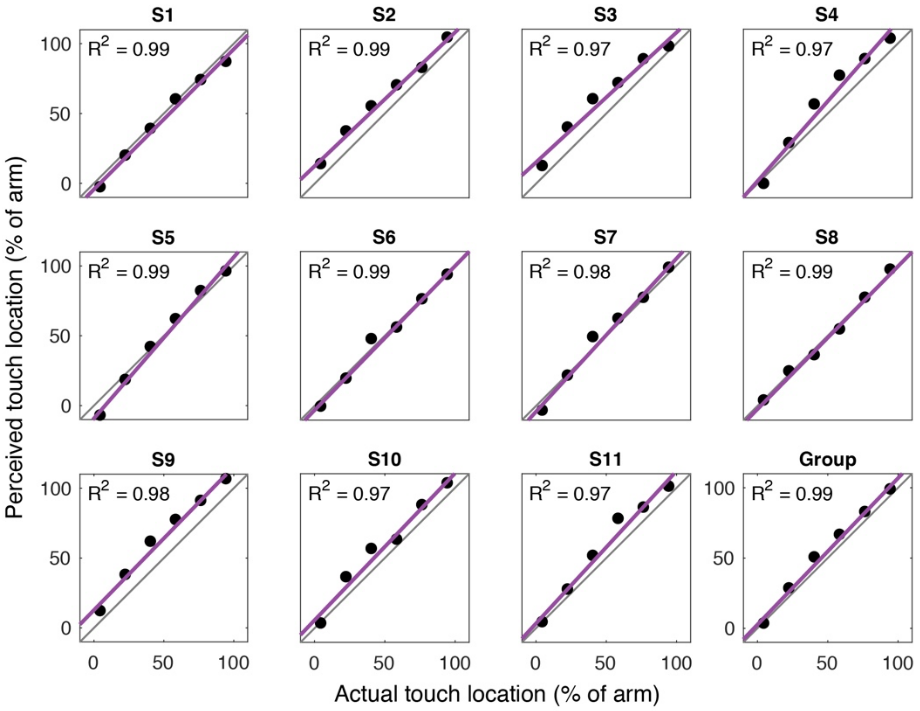

**Figure S3. Linear fits to localization for individual participants in Experiment 1**

The regression fits for each participant's localization and the corresponding goodness of fit. Each panel corresponds to an individual participant (S1–S11) or the group average (Group; bottom right). Gray line is the identity line (i.e., perfect localization).

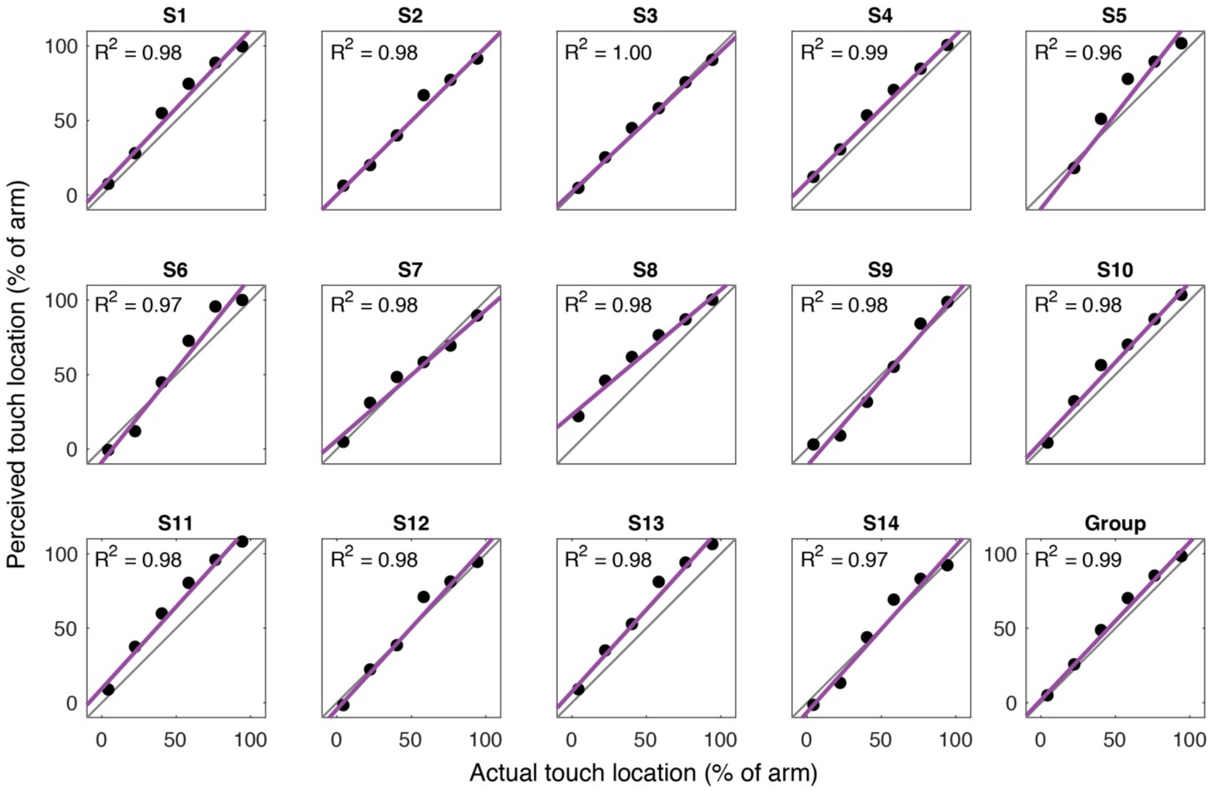

**Figure S4. Linear fits to localization for individual participants in Experiment 2**

The regression fits for each participant's localization and the corresponding goodness of fit. Each panel corresponds to an individual participant (S1–S14) or the group average (Group; bottom right). Gray line is the identity line (i.e., perfect localization).

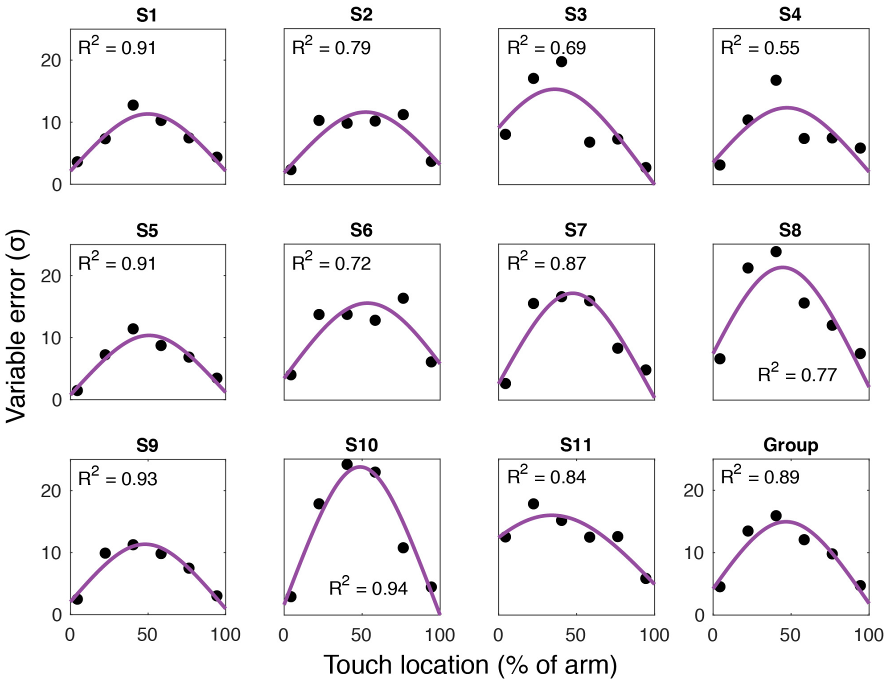

**Figure S5. Trilateration model fits for each participant's variable errors in Experiment 1**

The regression fits for each participant's variable errors (unit: % of surface) and the corresponding goodness of fit. Each panel corresponds to an individual participant (S1–S11) or the group average (Group; bottom right). As can be seen, our trilateration model provided a good fit to each participant's data.

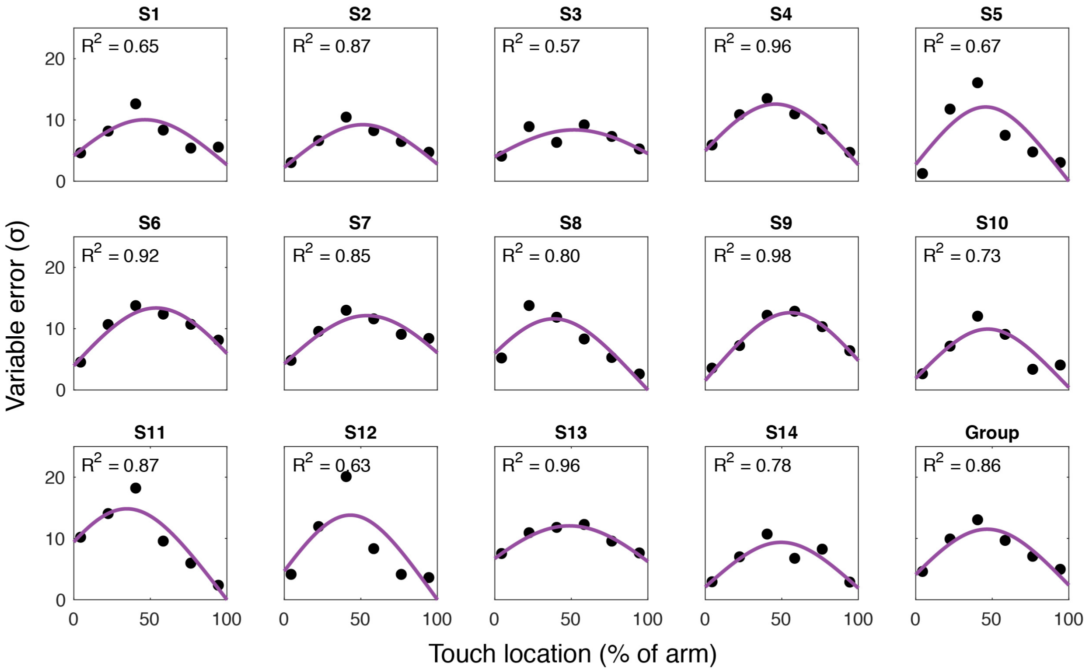

**Figure S6. Trilateration model fits for each participant's variable errors in Experiment 2**

The regression fits for each participant's variable errors (unit: % of surface) and the corresponding goodness of fit. Each panel corresponds to an individual participant (S1–S14) or the group average (Group; bottom right). As can be seen, our trilateration model provided a good fit to each participant's data.

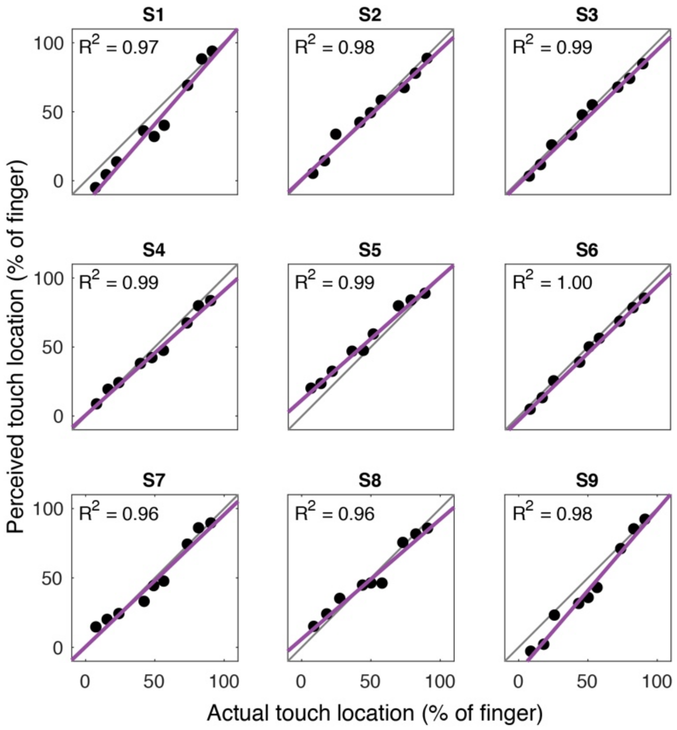

**Figure S7. Linear fits to localization for individual participants in Experiment 3**

The regression fits for each participant's localization and the corresponding goodness of fit. Each panel corresponds to an individual participant (S1–S9). Gray line is the identity line (i.e., perfect localization).

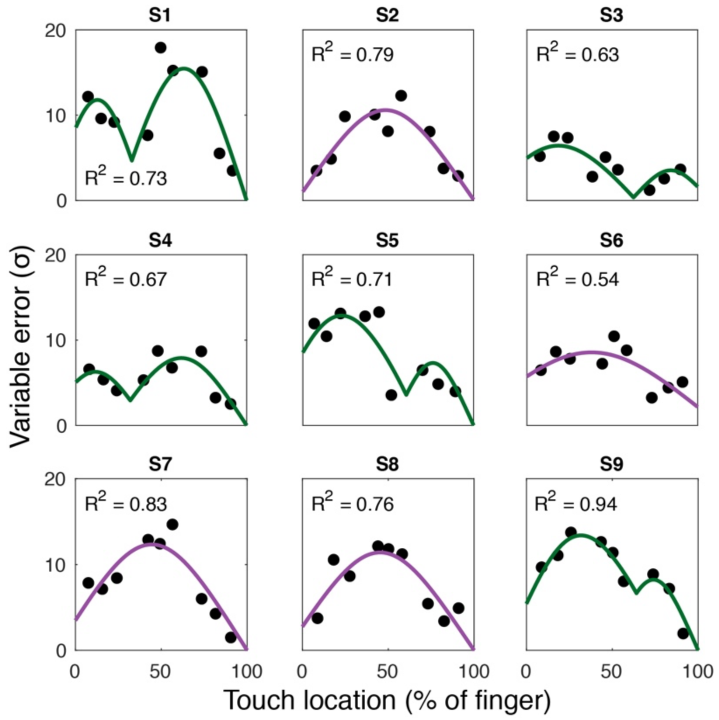

**Figure S8. Trilateration model fits for each participant's variable errors in Experiment 3**

The best fits for each participant's variable errors (unit: % of surface) and the corresponding goodness of fit. Each panel corresponds to an individual participant (S1–S9). As can be seen, our trilateration model provided a good fit to each participant's data. When the two-landmark model provided the best fit to the participant's data, the line is purple. When one of the three-landmark models provided the best fit to the participant's data, the line is green.

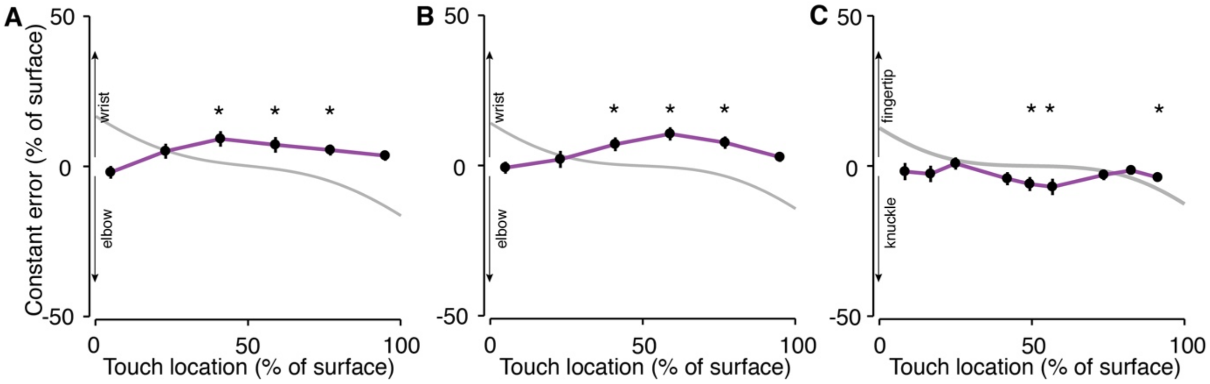

**Figure S9. Constant errors in both experiments**

The constant errors (% of surface) observed in both (A) Experiment 1 and (B) Experiment 2 demonstrate that localization was biased towards the wrist. The constant errors for (C) Experiment 3 were biased towards the knuckle. The gray line corresponds to the predicted pattern of error following boundary truncation, which was unable to fit the data. *$p$<.05 (uncorrected)

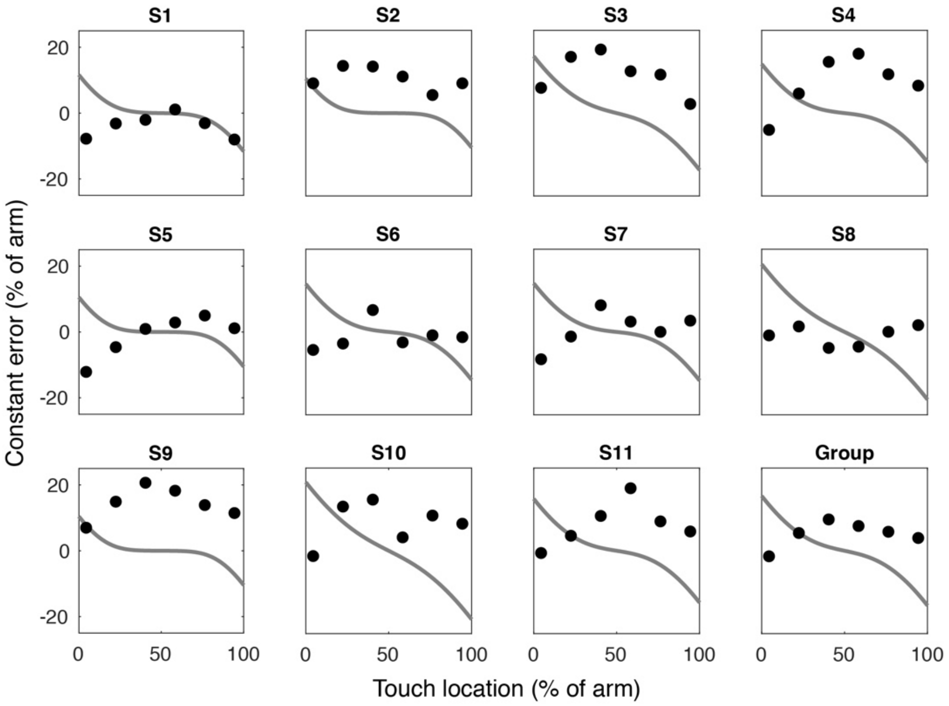

**Figure S10. Fits between constant error and boundary truncation in Experiment 1**

Here we show the relationship between the observed constant errors (% of arm) and the expected constant error bias given truncation at the boundaries of the arm (gray line). Each panel corresponds to an individual participant (S1–S11) or the group average (Group; bottom right). Boundary truncation provided a poor fit ($R^2$<0) for each participant.

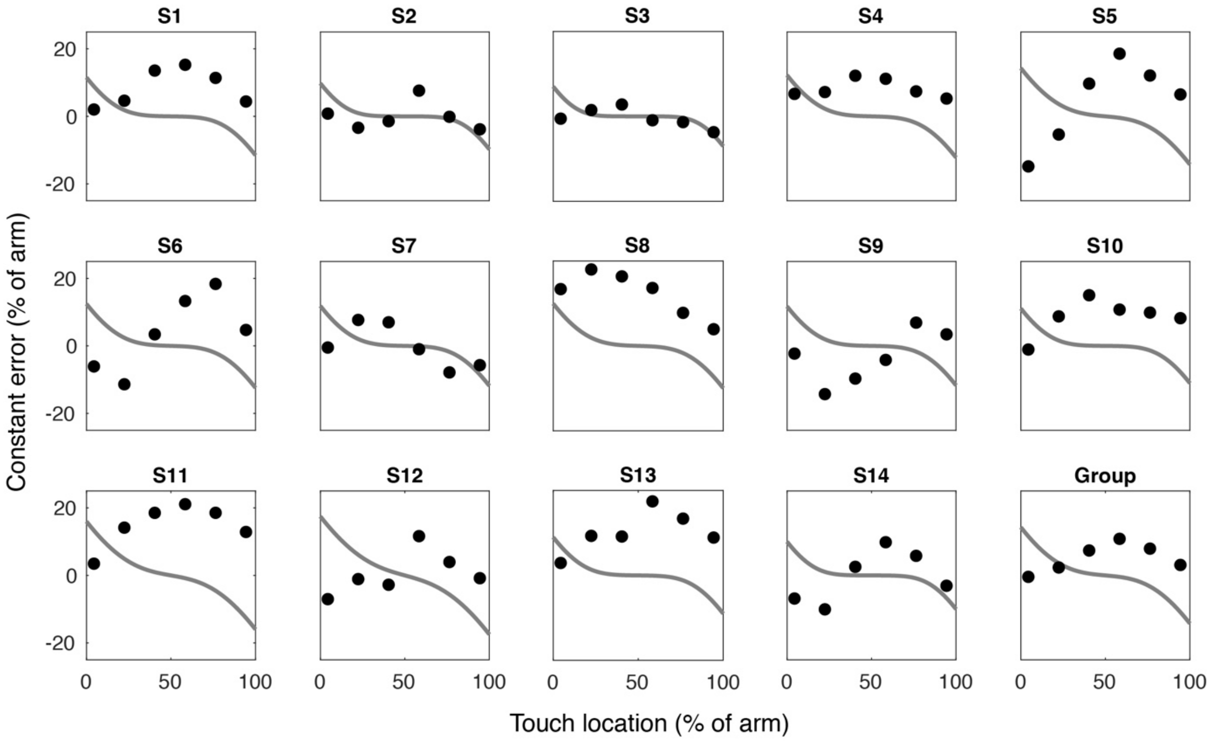

**Figure S11. Fits between constant error and boundary truncation in Experiment 2**

Here we show the relationship between the observed constant errors (% of arm) and the expected constant error bias given truncation at the boundaries of the arm (gray line). Each panel corresponds to an individual participant (S1–S14) or the group average (Group; bottom right). Boundary truncation provided a poor fit ($R^2$<0) for each participant.

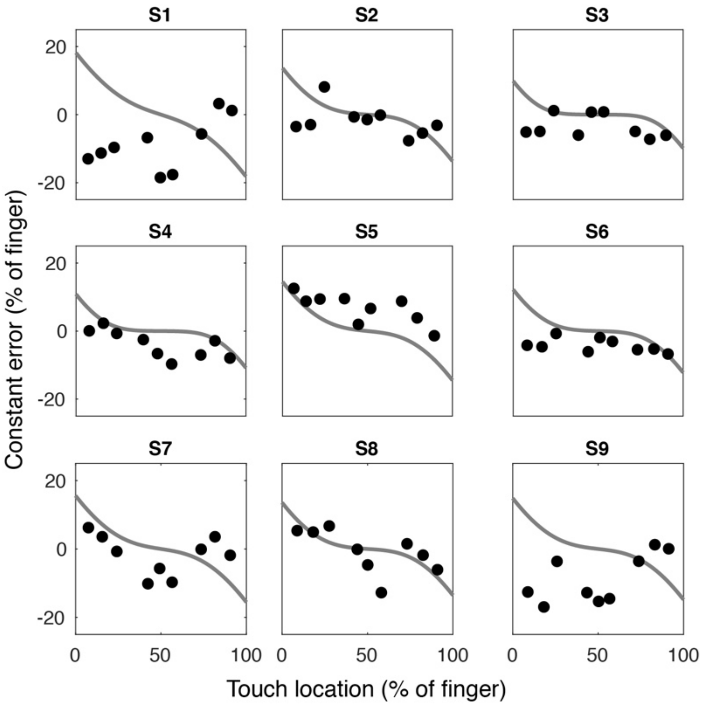

**Figure S12. Fits between constant error and boundary truncation in Experiment 3**

Here we show the relationship between the observed constant errors (% of finger) and the expected constant error bias given truncation at the boundaries of the finger (gray line). Each panel corresponds to an individual participant (S1–S9). Boundary truncation provided a poor fit ($R^2$<0) for each participant.

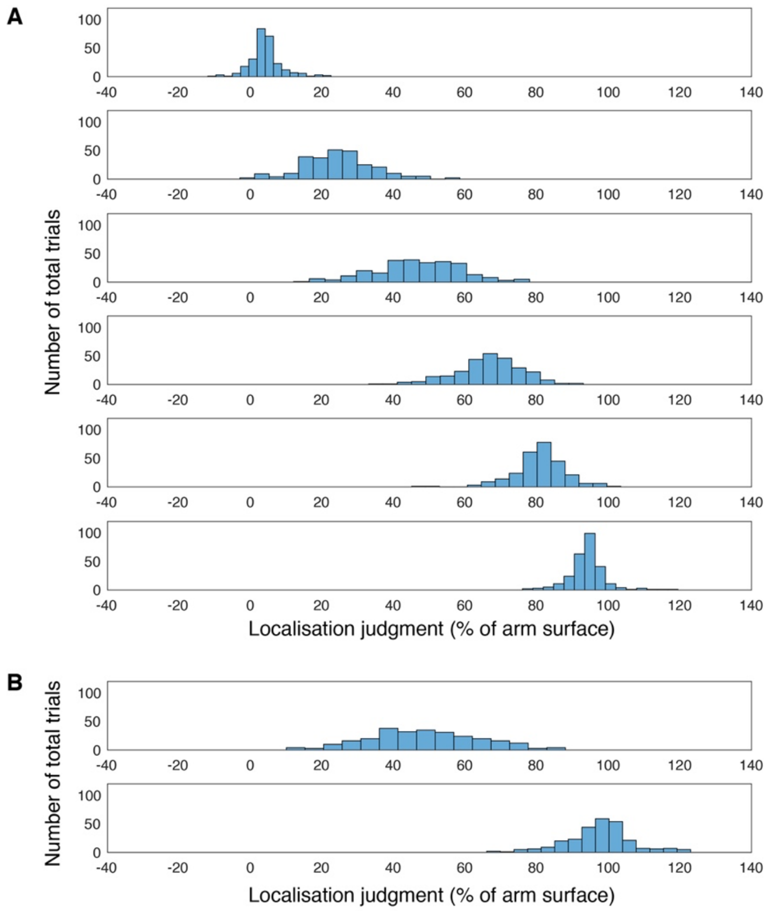

**Figure S13. No response truncation at the boundaries of the arm**

(A) Histograms showing the distribution of judgments (Exp. 2) across the arm (0%=elbow; 100%=wrist) observed at each location from closest to the elbow (top) to closest to the wrist (bottom). Each distribution has been centered on the group-level mean to remove participant-specific biases in localization. (B) Non-mean centered distribution of responses for the middle (top) and near-wrist touch locations (bottom) to illustrate the full spread of response location across the surface. Each location shows a bell-shaped distribution of responses. Furthermore, we did not observe any response truncation at the boundaries of the arm, as responses were made into the neighboring limbs (hand and lower arm). Therefore, our variable error results are not due to any truncation in the responses.

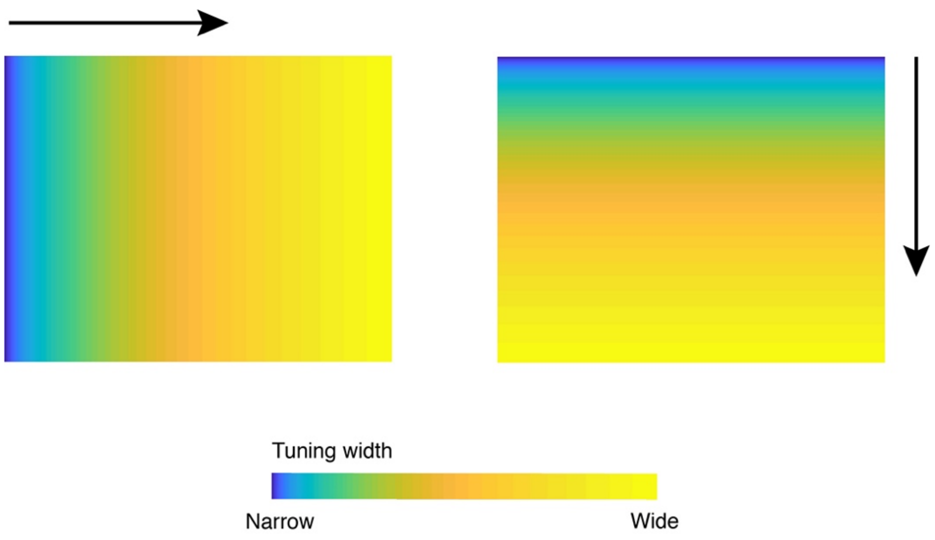

**Figure S14. Gradients in tuning width for one subpopulation in the 2D network**

Here we show the two-dimensional distance-dependent gradients in tuning width for one subpopulation. The gradient in the $x$-dimension is on the left and the $y$-dimension is on the right. The gradients in each dimension are independent of each other; each unit therefore multiplexes a distance combination in each dimension. Colors correspond to the tuning width (narrow-to-wide; blue-to-yellow) and the arrow denotes the direction of the gradient in the $xy$ plane.

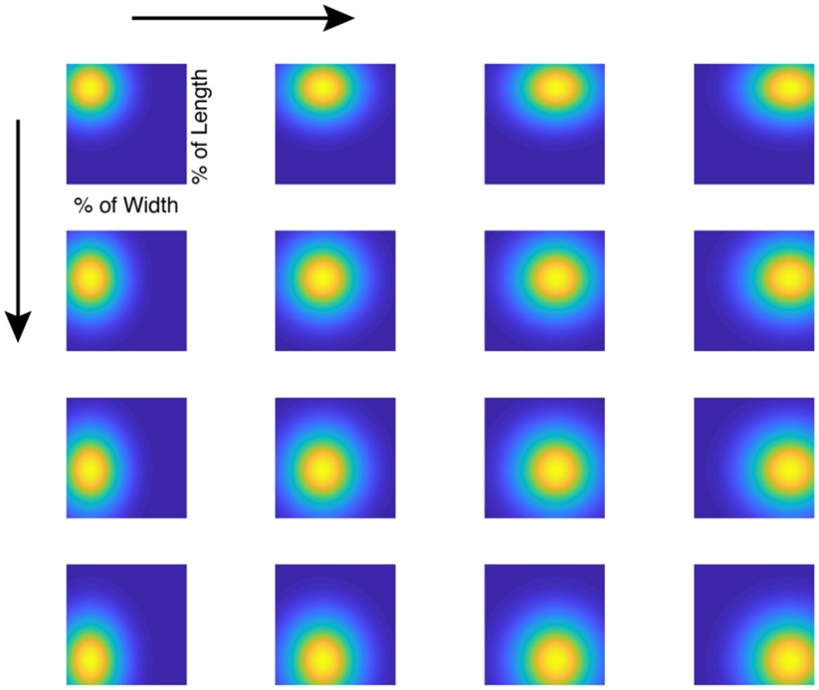

**Figure S15. Receptive fields in one subpopulation of the 2D network**

Receptive fields in one subpopulation (see, also Figure S14) tiling the entire $xy$ plane of the sensory surface. Each panel corresponds to the receptive field of an individual unit. The shapes of receptive fields in each subpopulation ranged from circular (upper left, bottom right) to oval-shaped. The width in each dimension of the receptive field reflects the distance from the landmark of that dimension. Arrows correspond to the direction of the gradient in tuning width.

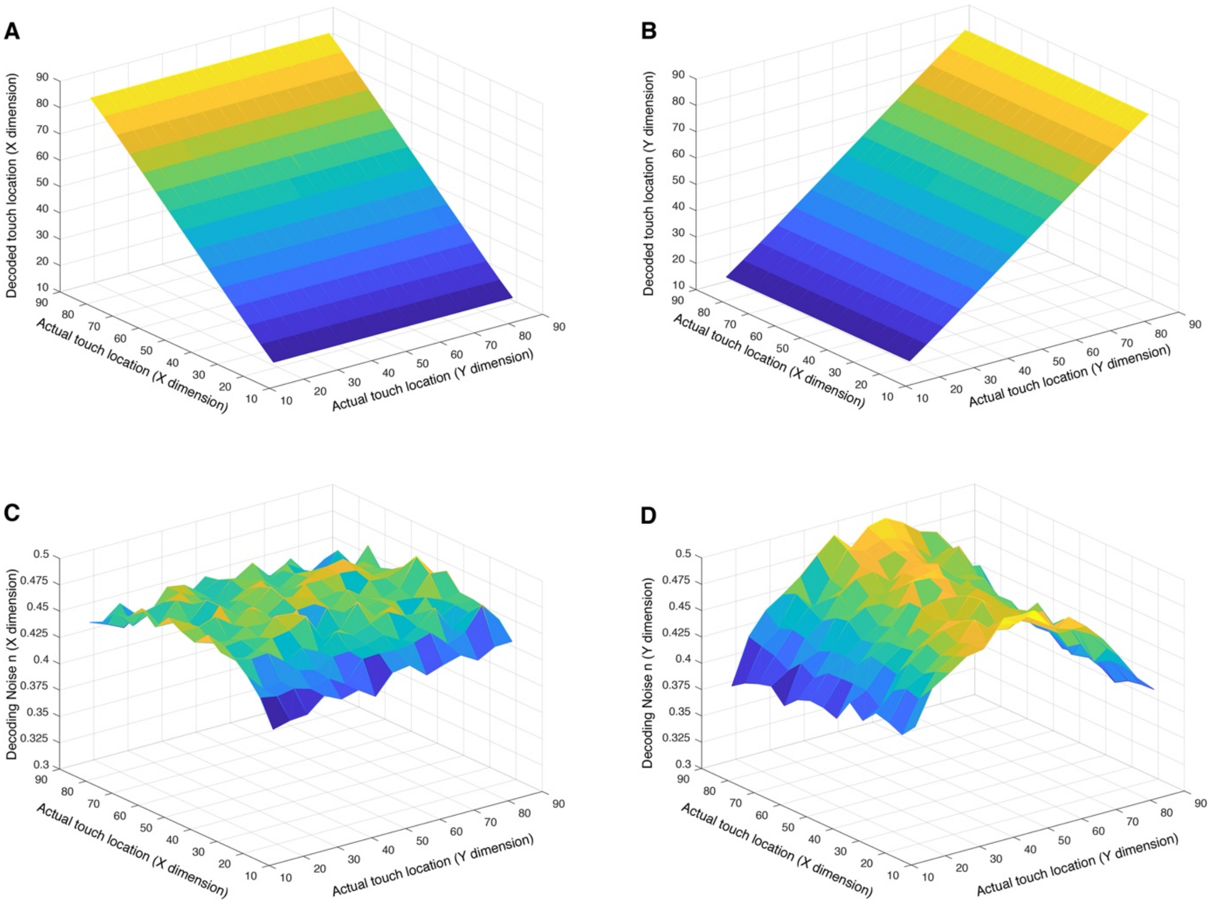

**Figure S16. Simulations results for the 2D network**

Localization in both the (A) $x$-dimension and the (B) $y$-dimension of the sensory surface was highly accurate. Importantly, trilateration with the multiplexed decoding populations produced inverted U-shaped patterns of variable errors. This was observed for both the (C) $x$-dimension and the (D) $y$-dimension of the sensory surface.

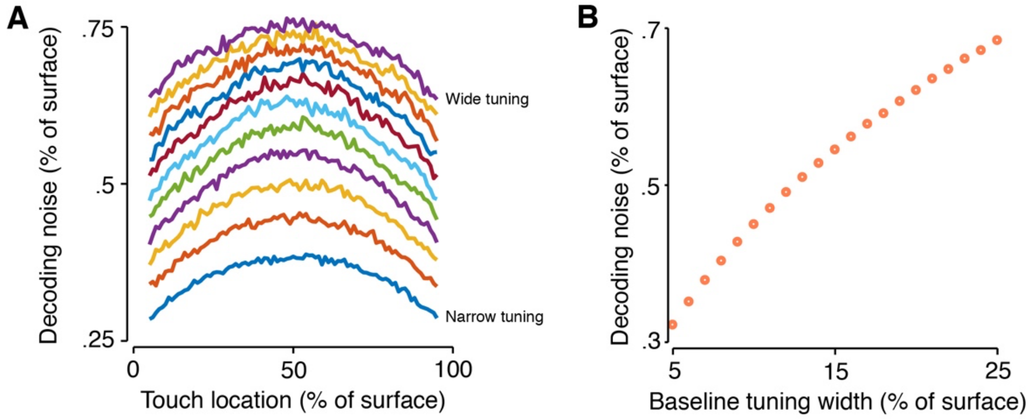

**Figure S17. Effect of tuning width on decoding noise**

The ability of our network to localize touch was partly dependent on the baseline tuning width (i.e., width at distance zero). As can be seen in (A), the offset of the inverted U-shaped pattern of decoding noise increased as tuning width increased. However, the magnitude and shape of the inverted U-shape (i.e., the range it covers) was virtually independent of baseline. The effect of offset can be seen in (B), where we show the mean decoding noise for each of the twenty-one baseline tuning values we simulated. The relationship between baseline tuning and mean decoding noise is almost linear. This can explain the well-known effect of receptive field shape and spatial precision. These stimulations also suggest that perceptual spatial precision has at least two factors: baseline tuning width and distance from a landmark.

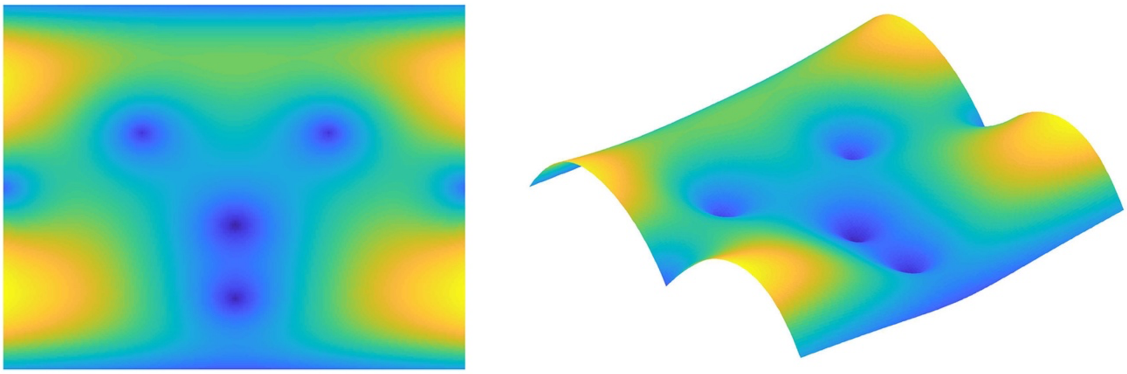

**Figure S18. Simulation results for tactile localization on the face**

Here we explored a possible pattern of variable error for tactile localization on a complex body part, the face. The face likely has a large diversity of landmarks that could contribute to localization. As with other body parts, we considered the boundaries of the face as landmarks (jawline and hairline). These were modelled as straight lines of distance-computing elements, consistent with the two-dimensional network described above. We also considered the mouth, the nose, the two eyes, and the two ears as landmarks. These landmarks were modelled as single points in face space. As can be seen above, tactile localization on the face is characterized by a complex pattern of variable error. The sides of the face reflect the inverted U-shaped pattern of variable error, whereas this pattern is "washed out" by the close proximity of multiple landmarks in the central portion of the face. Note that this is only a simulation; future work should investigate the true pattern of localization on the face.
